# The proteasome regulator PI31 is required for protein homeostasis, synapse maintenance and neuronal survival in mice

**DOI:** 10.1101/711515

**Authors:** Adi Minis, Jose Rodriguez, Avi Levin, Kai Liu, Eve-Ellen Govek, Mary E. Hatten, Hermann Steller

## Abstract

Proteasome-mediated degradation of intracellular proteins is essential for cell function and survival. The proteasome-binding protein PI31 (Proteasomal Inhibitor of 31kD) promotes 26S assembly and functions as an adapter for proteasome transport in axons. As localized protein synthesis and degradation is especially critical in neurons, we generated a conditional loss of PI31 in spinal motor neurons (MNs) and cerebellar Purkinje cells (PCs). A cKO of PI31 in these neurons caused axon degeneration, neuronal loss and progressive spinal and cerebellar neurological dysfunction. For both MNs and PCs, markers of proteotoxic stress preceded axonal degeneration and motor dysfunction, indicating a critical role for PI31 in neuronal homeostasis. The time course of the loss of MN and PC function in developing mouse CNS suggests a key role for PI31 in human developmental neurological disorders.

**Statement of Significance:** The conserved proteasome-binding protein PI31 serves as an adapter to couple proteasomes with cellular motors to mediate their transport to distal tips of neurons where protein breakdown occurs. We generated global and conditional PI31 knockout mouse strains and show that this protein is required for protein homeostasis, and that its conditional inactivation in neurons disrupts synaptic structures and long-term survival. This work establishes a critical role for PI31 and local protein degradation in the maintenance of neuronal architecture, circuitry and function. Because mutations that impair PI31 function cause neurodegenerative diseases in humans, reduced PI31 activity may contribute to age-related neurodegenerative diseases.

## Introduction

The regulated breakdown of proteins is essential for cell function and survival. The Ubiquitin-Proteasome System (UPS) is responsible for the degradation of soluble intracellular proteins, whereas autophagy primarily serves to remove protein aggregates and damaged organelles (1–8). UPS-mediated protein degradation involves two major steps: first, a protein is tagged with a chain of ubiquitin molecules, and then poly-ubiquitinated substrates are cleaved into small peptides by the 26S proteasome (2, 9, 10). Although protein homeostasis is thought to be a critical feature of neuronal circuit function, the major emphasis has been the importance of local protein translation (11, 12). It has been suggested that local protein degradation in distal axons and dendrites is important for quality control of synaptic proteins, and also for plasticity and remodeling of synapses (13–20). However, in the absence of tools to selectively inhibit proteasomes at synapses the precise role of local proteasome activity remains unclear.

Proteasome activity is regulated at multiple levels, including transcription of proteasomal proteins, assembly of subunits, post-translational modifications, and the utilization of different regulatory particles (10, 21–23). One structurally conserved proteasome regulator is PI31 (“<u>p</u>roteasome inhibitor of <u>31</u>kD”). PI31 was originally discovered based on its ability to inhibit hydrolysis of small peptides by 20S particles *in vitro* (24, 25). Conversely, studies in *Drosophila*, yeast and plants indicate that PI31 promotes protein breakdown *in vivo* (26–28). PI31 can be ADP-ribosylated by Tankyrase, and this modification promotes assembly of 26S proteasome (29). PI31 also binds directly to Fbxo7/PARK15, a conserved F-box protein that is the substrate-recognition component of an SCF E3-ligase complex (26, 30–32). Unexpectedly, loss of Fbxo7 causes cleavage and inactivation of PI31, reduced proteasome activity and neuronal degeneration (26, 33, 34). Significantly, impaired UPS function and mutations in Fbxo7/PARK15 are associated with neurodegenerative diseases (33–42).

We recently showed that in addition to its effect on proteasome assembly, PI31 is an adapter for neuronal proteasome transport, suggesting a key role in protein homeostasis and synaptic function (43). To examine the physiological function of PI31, we generated global and conditional knockout mouse strains and investigated how loss of PI31 affects two major types of neurons - spinal motor neurons (MNs) and cerebellar Purkinje cells (PCs). Spinal MNs reside in the ventral horn of the spinal cord, while their long axons, that can extend over one meter in humans, innervate effector muscles at a specialized synapse called the neuromuscular junction (NMJ) (44). PCs are the sole output neurons of the cerebellar cortex. Their dendrites receive inputs from cerebellar granule cell parallel fibers and inferior olivary nucleus climbing fibers, while their axons project through the inner granular layer (IGL) of the cerebellar cortex, where mature granule cells (GCs) and Golgi interneurons reside, to synapse onto deep cerebellar nuclei neurons (DCNn) in the deep cerebellar nuclei (DCN) (45, 46). Both of these neurons are also involved in a wide range of neurodegenerative diseases, including Amyotrophic Lateral Sclerosis (ALS) and Spinal Muscular Atrophy (SMA) in the case of spinal MNs, and ataxias, autism and cerebellar essential tremor in the case of PCs (47–53).

Our study reveals that PI31 is an essential gene, as knockout embryos died at mid to late-gestation, although the development and differentiation of many embryonic cell types appeared normal overall. Strikingly, in both motor neurons and Purkinje cells of the cerebellum, loss of PI31 function impaired protein homeostasis in neuronal processes, disrupted the architecture of synapses, axons and dendrites, and compromised motor function. Importantly, degeneration of neuronal synapses and processes became progressively more severe with age, culminating in neuronal loss. Inactivation of PI31 in MNs and PCs recapitulated the progressive neuropathology and motor dysfunction of previously described mouse models of ALS and ataxia, respectively, and was reminiscent of the severe behavioral and anatomical defects associated with human spinal MN and PC neurodegenerative diseases (54–58). Collectively, this work establishes a critical role for PI31 and protein degradation in the maintenance of neuronal architecture, circuitry and function. Because mutations that impair PI31 function are thought to cause neurodegenerative diseases in humans, reduced PI31 activity may contribute to age-related neurodegenerative diseases.

## Results

### Generation of PI31-Null mice

In order to examine the physiological role of PI31, we used two independent approaches to generate constitutive and conditional PI31 loss-of-function mouse mutants (Fig. 1). First, we used CRISPR/Cas9 technology to introduce a 16 bp deletion in exon 1 of the mouse PI31 gene. This resulted in a frame shift and a premature stop codon (Fig. 1A, B, C). Western blot analysis confirmed that no PI31 protein was detectable in PI31^CRISP/CRISP^ embryos, indicating that this allele is a Null-mutant (Fig. 1D). PI31^CRISP/CRISP^ embryos appeared normal overall prior to embryonic stage E13.5. However, at E15.5 small size differences became detectable and by E18.5 reduced growth was apparent (Fig. 1E). Histological analysis of these embryos indicated that many tissues and organs, such as heart, kidney, gut, muscle and skin, developed normally overall and contained differentiated cells with appropriate morphology and patterning (Fig. S1). We conclude that PI31 function, unlike complete inactivation of proteasomes, is not universally required for the growth, differentiation and survival of many cell types.

**Fig. 1:**
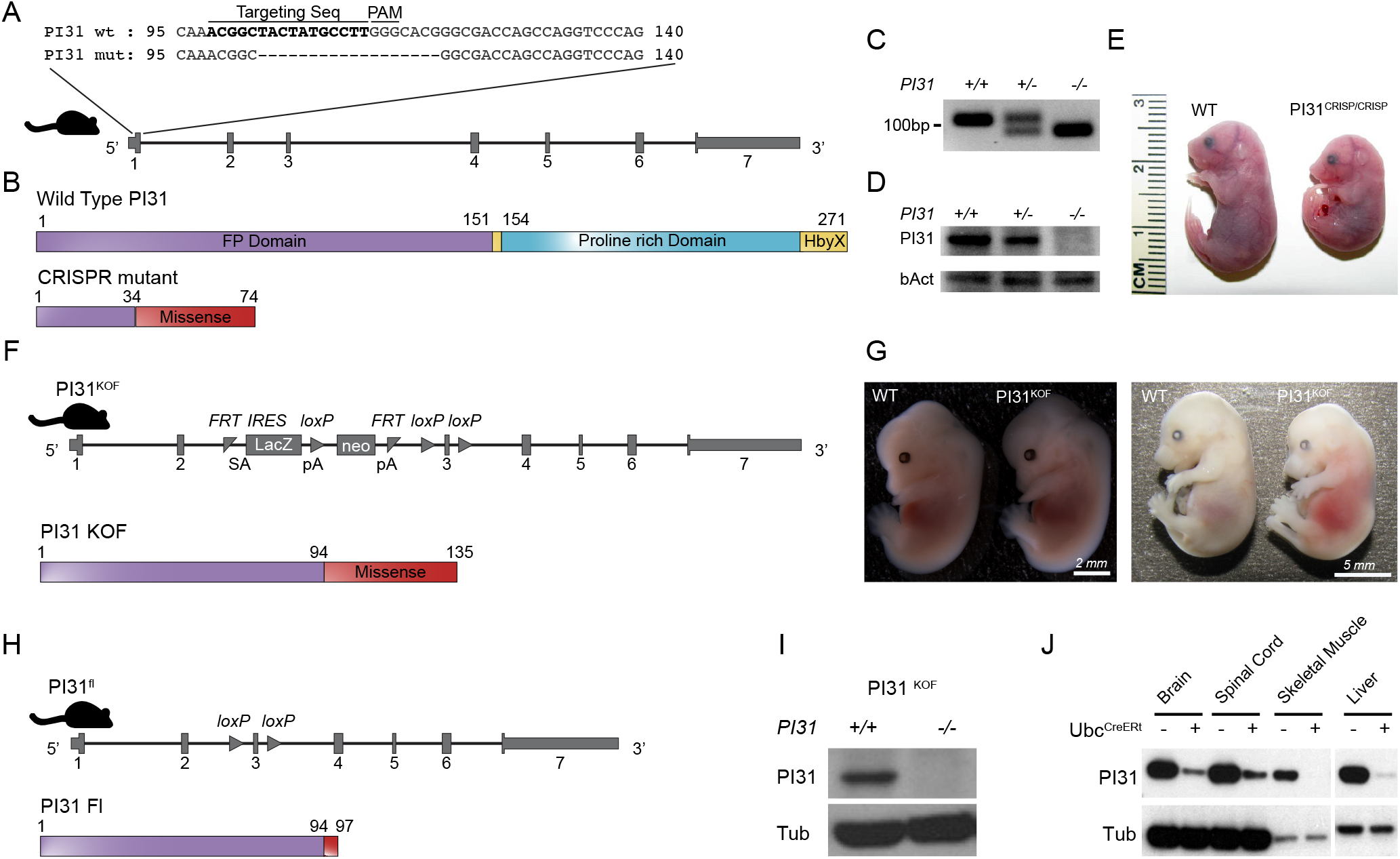
Inactivation of PI31 results in embryonic lethality. Two independent approaches were used to generate constitutive and conditional PI31 loss-of-function mouse mutants. **A.** CRISPR/Cas9 technology was used to introduce a 16 bp deletion in exon 1 of the mouse PI31 gene. Alignment of WT and PI31^CRISP^ mutant DNA sequences illustrates the extent of this deletion; the sequence of the targeting gRNA is depicted in bold. The deletion causes a frame shift and generates a premature stop codon. **B.** Schematic presentation of wild type PI31 and PI31^CRISP^ protein domains. The mutant protein derived from the PI31^CRISP^ allele is predicted to retain the first 34 amino acids (encoded by nucleotides 1-102 of the PI31 coding sequence), followed by 40 missense amino acids and a premature stop codon. (FP domain – required for homo- and hetero-dimerization with other PI31 molecules and\or the E3 ligase FBXO7, Proline Rich Domain – proline rich unstructured domain, HbyX domain – a domain commonly found in modulators of proteasome activity and considered to be important for binding the 20S proteasome particle). **C.** PCR analysis of WT and PI31^CRISP^ DNA confirms the presence of a deletion in the PI31^CRISP^ mutant. **D.** PI31 protein levels were examined by Western blot analysis of brain extracts from E13.5 embryos. PI31 protein was readily detected in wild type embryos, but levels were reduced in PI31^CRISP/+^ and not detectable in PI31^CRISP/CRISP^ embryos. **E**. PI31^CRISP/CRISP^ mutants are embryonic lethal. Representative pictures of WT and PI31^CRISP/CRISP^ embryos at E18.5 indicate growth retardation and developmental delay. **F.** Illustration of the gene architecture and predicted protein product in the knockout first (KOF) PI31^KOF^ allele. A PI31 knockout mouse mutant strain (PI31^KOF^) was generated from commercially available ES cell (Psmf1^tm1a(EUCOMM)Hmgu^). LacZ is translated from internal ribosome entry site (IRES), and this insertion is predicted to truncate PI31 by retaining the first 94aa encoded in exons 1 and 2, followed by 41 missense amino acids and a stop codon. **G.** PI31^KOF^ homozygotes are embryonic lethal. Representative pictures of WT and PI31^KOF/KOF^ embryos at ages E13.5 and E15.5. **H.** PI31^KOF^ mice were bred with a strain expressing FLP1 recombinase in order to generate a PI31 allele where the third exon is flanked by two loxP sites (PI31^fl^, Psmf1^tm1c(EUCOMM)Hmgu^). After Cre recombination, the mutant protein derived from the PI31^fl^ allele is predicted to retain the first 94 amino acids, followed by 3 mutant amino acids and a stop codon. **I,J**. Validation of PI31 KO by western blot. **I.** No PI31 protein was detectable in MEFs derived from P31^KOF/KOF^ embryos. **J.** PI31^f/f^ mice were bred with mice expressing a Tamoxifen-inducible Cre recombinase under control of the Ubc promoter. Reduced PI31 protein levels were measured in multiple tissues 2 weeks after tamoxifen injection.

Because PI31 complete knockout mutants died at advanced embryonic stages, we generated conditional PI31 mutants in order to more carefully examine the role of PI31 in post-embryonic cell types. For this purpose, we used the “knockout-first” (KOF) strategy which allows the production of reporter knockouts, conditional knockouts, and Null-alleles following exposure to the sitespecific recombinases Cre and Flp (59, 60). A PI31 knockout-first (PI31^KOF^) mutant line was generated from commercially available ES cell (Psmf1^tm1a(EUCOMM)Hmgu^) as illustrated in Fig. 1F. The lacZ insertion introduces a strong splice acceptor, which is predicted to cause severe truncation of the PI31 protein and eliminate domains critical for function (Fig. 1F). Similar to PI31^CRISP/CRISP^ mice, PI31^KOF/KOF^ mice are recessive embryonic lethal, and by E15.5 embryos were hemorrhagic and smaller in size (Fig. 1G). Western blot analysis confirmed that PI31 protein was not detectable in PI31^KOF/KOF^ embryos (Fig. 1I). We conclude that both the PI31^CRISP/CRISP^ and PI31^KOF/KOF^ strains represent loss-of-function mutants in which expression of PI31 protein was successfully ablated, and that PI31 is an essential gene for mouse development. In order to generate conditional alleles for more in-depth analysis of PI31 function in specific post-embryonic cell types, PI31^KOF^ mice were bred with mice expressing FLP1 recombinase to generate a PI31 allele where the third exon is flanked by two loxP sites (PI31^fl^, Psmf1^tm1c(EUCOMM)Hmgu^)(Fig. 1H). PI31^fl/fl^ mice express normal levels of PI31, are viable, fertile and appear completely normal. However, introduction of various Cre-drivers successfully inactivated PI31 and led to tissue specific loss of PI31 protein, as indicated in Fig. 1J.

### Inactivation of PI31 causes neurological defects, proteotoxic stress and reduced proteasome assembly

We used PI31^fl/fl^ mice to investigate the physiological role of PI31 in different types of neurons. First, we crossed PI31^fl/fl^ mice to CDX2-Cre mice to inactivate PI31 in the caudal part of the embryo. PI31^fl/fl^/CDX2-Cre mice were viable but around P6 they began to develop progressive neuromotor phenotypes, characterized by spasticity, rigid muscle tone, strong tremor and a severely impaired righting response (Fig. 2A, Movie S1). When these mice were picked up by their tail, they displayed hind leg clasping between episodes of tremor, a hallmark of neuromotor dysfunction (Fig 2A). Mutant mice were only able to move using their front legs since their hind limbs were hyperextended and paralyzed (Fig. 2A, Movie S1). These phenotypes became progressively more severe with age, and by 3-4 weeks all mice succumbed. We also noted that although PI31 was deleted in all caudal tissues – including skin, muscle and kidney – we did not find any significant phenotype in these tissues.

**Fig. 2:**
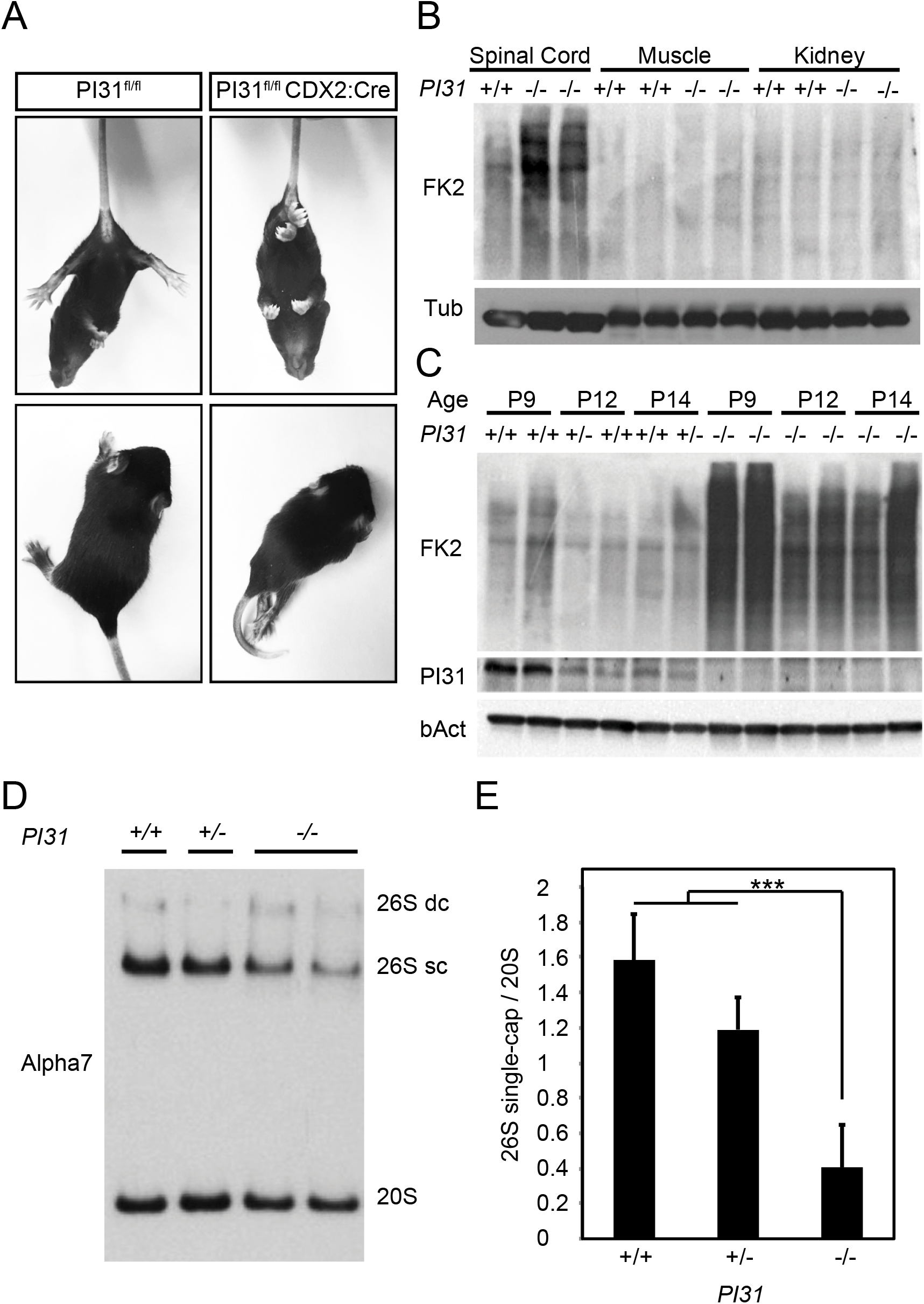
Caudal inactivation of PI31 leads to severe motor defects and proteotoxic stress in the spinal cord. **A.** Representative pictures of PI31^fl/fl^ CDX2:Cre mice and a control littermate 14 days postnatal. The upper two panels depict hind feet clasping, and two lower panels illustrate hind limb paralysis in mutant mice. **B.** Western blot analysis of spinal cord, muscle and kidney of PI31^fl/fl^ CDX2:Cre mice and their control littermates at P14. Blotting for poly-ubiquitinated proteins (FK2) indicates that spinal cord tissue was more sensitive to the loss of PI31 than muscle or kidney. **C.** Western blot analysis of spinal cord extracts from PI31^fl/fl^ CDX2:Cre mice and control littermates at indicated ages. Immuno-blots for ubiquitin revealed a dramatic accumulation of poly-ubiquitinated proteins in PI31 mutants. PI31 expression in wild type spinal cords was correlated with the accumulation of poly-ubiquitinated proteins in age-matched PI31 knockouts. **D.** Mouse embryonic fibroblasts (MEF) derived from PI31^KOF^ embryos have decreased levels of 26S proteasomes. Western blot analysis following native gel electrophoresis of WT, heterozygous and PI31 knockout MEFs revealed a reduction in the levels of single-capped 26S proteasome. 26S double-capped (26S dc), single capped (26S sc) and 20S proteasomes were detected with an anti-Alpha7 proteasome subunit antibody. **E.** Quantification of proteasome profiles. Densitometry analysis of single capped 26S and 20S proteasomes was performed, and the ratio of single capped 26S to 20S proteasomes was calculated for each sample. Pooled data of densitometry analysis of 3 independent experiments show a reduction of single capped 26S proteasomes in PI31 knockout MEFs. Statistical analysis was performed using one-way ANOVA test, *** stands for p-value = 0.001.

Next, we looked for evidence that protein homoestasis was disrupted in PI31^fl/fl^/CDX2-Cre mice. For this purpose, we used the FK2 antibody to detect accumulation of poly-Ub proteins, which serves as a readout for proteasome function (61). While no differences between control and PI31^fl/fl^/CDX2-Cre were seen in protein extracted from muscle or kidney, we detected a clear and significant accumulation of poly-Ub proteins in spinal cord extracts from mutant mice (Fig. 2B). Highest levels of FK2 staining were seen in P9 mutant mice, at a time when motoric problems were apparent, but well before the peak of this phenotype and death of mutant animals (Fig. 2C). These observations indicate that inactivation of PI31 in the spinal cord leads to defects in the degradation of poly-Ub proteins, consistent with impaired proteasome function. It was previously shown that PI31 can promote the assembly of active 26S proteasomes from 19S and 20S particles (29). Therefore, we investigated whether loss of PI31 affects the proteasome profile. For this purpose, we used Western blot analysis of native gels to analyze extracts from mouse embryonic fibroblasts (MEF) derived from PI31^KOF/KOF^ embryos and controls (Fig. 2D, E). This revealed a small but significant reduction in the levels of single-capped 26S proteasomes, without substantial change of individual subunits (Fig. S2A). It is worth noting that PI31-Null MEFs were viable for extended periods in culture, indicating that these biochemical changes and the modest accumulation of poly-Ub conjugates in mutant MEFs under normal growth conditions did not have profound biological consequences (Fig. S2). We also saw induction of p62, which is a hallmark of proteotoxic stress (Fig. S2B), (62, 63). Since p62 delivers poly-Ub aggregates to the autophagic machinery for degradation in lysosome, it is possible that this backup pathway can clear proteins escaping degradation in PI31-Null MEFs sufficiently well enough to avoid overt biological consequences. We also treated MEFs with different concentrations of the proteasome inhibitors bortezomib and MG132 to look for a role of PI31 under conditions where protein breakdown is compromised (64). For both proteasome inhibitors, we observed that PI31-Null MEFs survived less well (Fig. S2C, D). This reveals a modest but significant requirement for PI31 function in fibroblasts under stress conditions. Taken together, our results indicate that loss of PI31 causes decreased proteasome activity and reduces the breakdown of poly-Ub proteins, but that this has little biological consequence for many cell types. In contrast, neurons appear to be particularly sensitive to the loss of PI31 function.

### Inactivation of PI31 in motor neurons disrupts the neuro-muscular junction (NMJ) and models ALS phenotypes

In order to examine a possible requirement of PI31 in neurons in more detail, we used several specific neuronal Cre-lines. To ablate PI31 function specifically in motor neurons we used Hb9-Cre (65, 66). PI31^fl/fl^ Hb9-Cre mice were initially viable but developed motor defects, kyphosis of the spine and muscle atrophy by the age of 5 months that became progressively more severe with age (Fig. 3 A, B). External inspection of these mice revealed severe kyphosis due to atrophy and weakness of paraspinal muscles, and also overt motoric defects that recapitulate phenotypes described for mouse ALS models (Fig. 3A, Movie S2) (53, 54, 67). Histological analysis of thoracic cross sections of 5-month old PI31^fl/fl^ Hb9^cre^ mice revealed highly atrophied musculature in PI31 mutants (Fig. 3A) and concomitant reduction in body weight (Fig. 3B). Next, we examined the architecture of the Neuro-Muscular Junction (NMJ) by staining motor neuron axons with β3-Tubullin/Synapsin, and the post-synaptic muscle end plate with α-Bungarotoxin (Fig. 3C). Compared to age-matched controls, PI31 mutants displayed severe structural abnormalities, including presynaptic fragmentation of the NMJ, axonal tip swellings, and massive axonal sprouting that are all notable at the age of 5 months (Fig. 3C). These phenotypes recapitulate defects described for several other mouse ALS models (53, 67, 68). The swelling of axonal tips was apparent in PI31-Null motor neurons as early as 1 month after birth and became progressively more severe with age (Fig. 3D). Interestingly, no overt morphological abnormalities were detectable in motor neuron cell bodies at this stage (data not shown). Moreover, conditional inactivation of PI31 in muscle did not cause any striking anatomical or behavioral phenotypes, even in mice that were aged up to 8 months. Finally, we also observed accumulation of p62-positive aggregates at the NMJ of PI31^fl/fl^ Hb9-Cre mice, but not in control littermates, indicating that loss of PI31 disrupts proteostasis in axon terminals of motor neurons (Fig. 3E). Notably, these phenotypes are remarkably similar to those caused by conditional inactivation of Rpt3, a critical component of the proteasome regulatory particle (54, 69). Rpt3, also termed PSMC4, is a critical component of the proteasome regulatory particle and required for the degradation of most proteasome substrates (69).

**Fig. 3:**
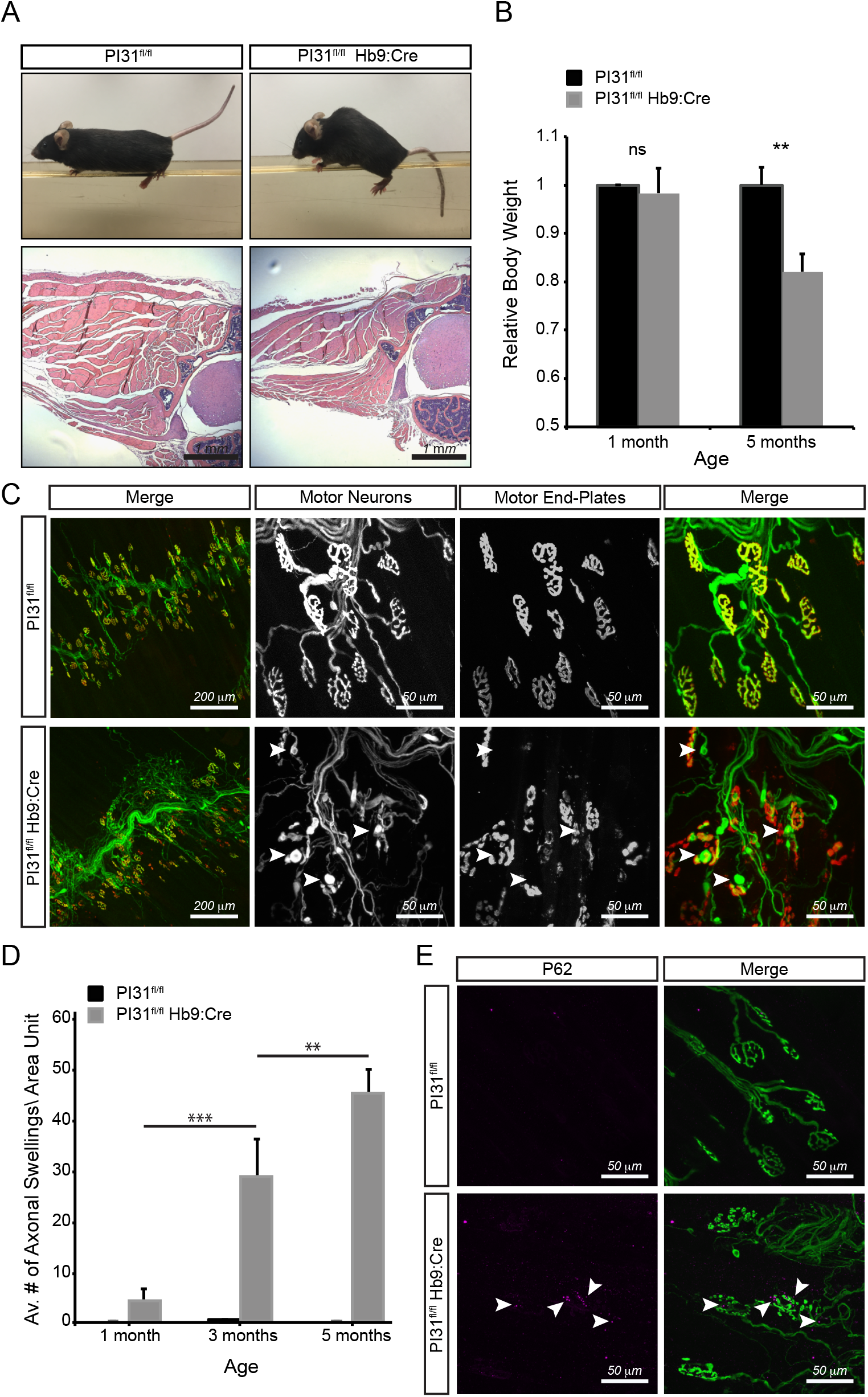
Inactivation of PI31 in motor neurons causes progressive muscle denervation and atrophy. **A.** PI31^fl/fl^ Hb9:Cre mice develop motor defects, kyphosis of the spine and muscle atrophy. Representative picture of PI31^fl/fl^ Hb9:Cre mice and their WT controls at 6 months of age. A representative H&E staining of thoracic cross sections of 5-month old PI31^fl/fl^ Hb9^cre^ mice and control littermates (PI31^fl/fl^ without Cre) showing atrophied musculature of KO mice. **B.** Relative body weight of 1 and 5 months old PI31^fl/fl^ Hb9:Cre mice and their controls littermate. PI31^fl/fl^ Hb9:Cre mice suffer progressive weight loss. 1MO – control, n=8, KO=4, 5MO - control, n=8, KO=4. Weight of WT controls at each age was set to 1. Statistical analysis was performed with a two-tailed paired t test, ** stands for p-value < 0.01. **C**. Inactivation of PI31 in motor neurons disrupts the structure of the Neuro-Muscular Junction (NMJ). Innervation of the Triangularis Sterni muscle was visualized by staining for □3-Tubullin/Synapsin (green), and post-synaptic muscle end plates were visualized with □-Bungarotoxin (red). 5 month old PI31^fl/fl^ Hb9:Cre mutant mice have fragmented NMJs, axonal swellings (indicated with white arrowheads) and massive axonal sprouting **D**. Loss of PI31 in motor neurons results in a progressive pathology, illustrated by the increase in axonal tip swellings with age. A bar diagram of the average number of axonal swelling per area unit (AU = 850 m^2^) at different ages as annotated. ** stands for p-value < 0.01, *** stands for p-value < 0.001 **E.** P62 granules accumulate at the NMJ of PI31^fl/fl^ Hb9:Cre, but not in control littermates. Representative image of NMJ in the Triangularis Sterni muscle of PI31^fl/fl^ and PI31^fl/fl^ Hb9:Cre mice (4MO females). P62 granules in magenta (marked with white arrowheads), motor neurons in green (stained by anti □3-Tubullin and Synapsin antibodies).

### Inactivation of PI31 in Purkinje cells causes progressive motor defects and disrupts protein homeostasis

We also examined the role of PI31 in the cerebellum, as it provides an outstanding model for studying how proteotoxic stress impairs neuronal function, structure and survival (70–78). Purkinje cells are very large neurons that are particularly sensitive to proteotoxic stress, which is thought to be a major factor contributing to cerebellar diseases (71, 79). As the Purkinje cell is the sole output neuron of the cerebellar cortex, defects in Purkinje cell function cause both motor and cognitive dysfunction. We used the Purkinje cell-specific Cre driver *Tg*(*Pcp2-Cre*) to examine how conditional loss of PI31 affects the health, structure, and viability of these cells as well as associated mouse behaviors. Behavioral analysis of *PI31^fl/fl^ PCP2-cre* mice revealed a series of defects characteristic for impaired Purkinje cell function, and these defects became progressively more severe with age. First, we used the ledge test and gait analysis to look for possible abnormalities in motor coordination in *PI31^fl/fl^ Pcp2-Cre* mice (Fig. 4B-D). At P22, *PI31^fl/fl^ Pcp2-Cre* mice were indistinguishable from their *PI31^fl/fl^* littermates. However, by P30, *PI31^fl/fl^ Pcp2-Cre* mice displayed an aberrant halting gait and frequently lost their balance on the ledge test (Fig. 4B). By P60, mutants had severely disrupted balance, frequently dragged their bodies along the ledge and fell on their heads when lowering themselves into the cage. Locomotor defects were also seen with gait analysis in older mutant mice (Fig. 4C; Movie S3). Taken together, these behavioral abnormalities suggest defects in Purkinje cell function.

**Fig. 4:**
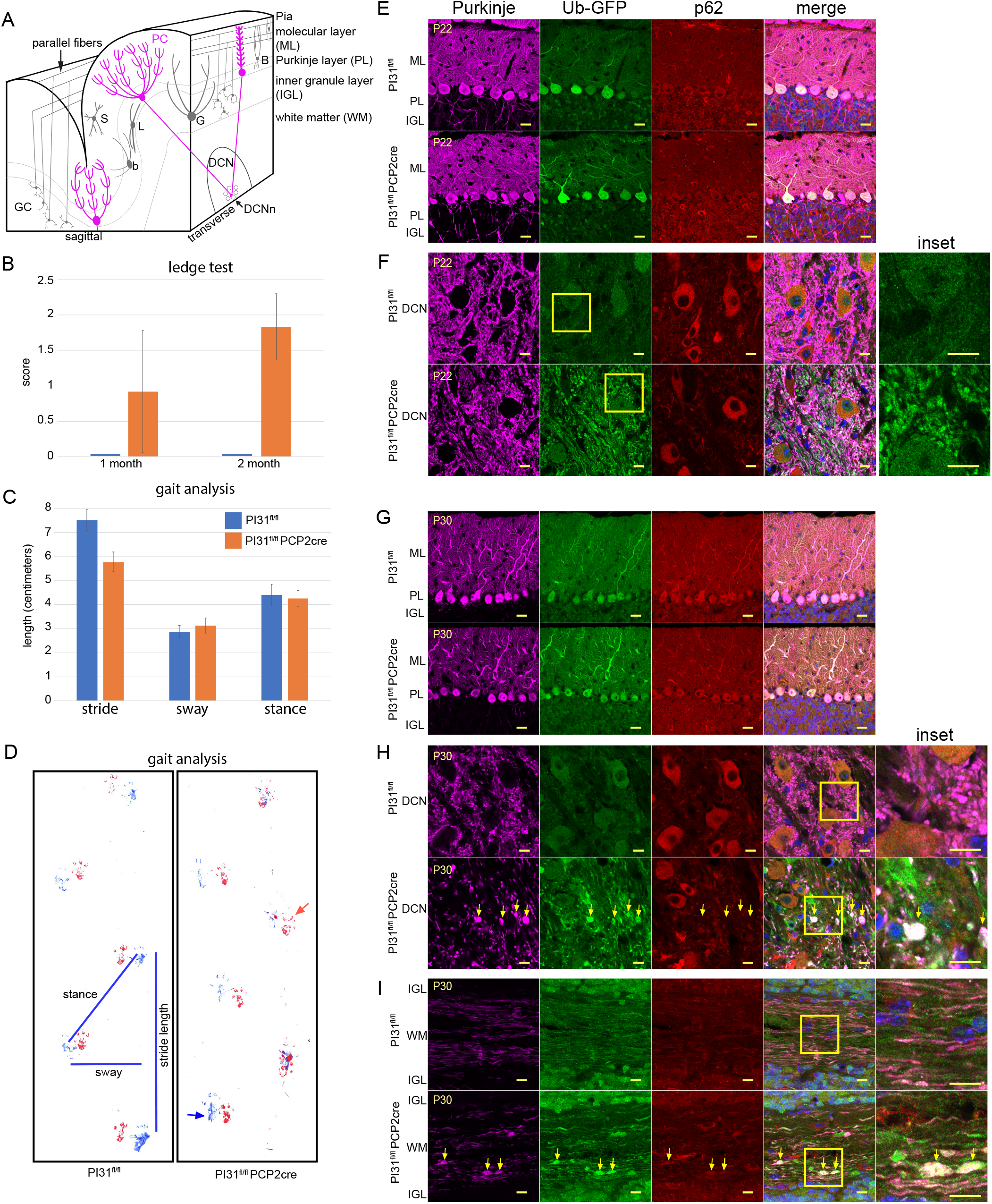
Loss of PI31 in Purkinje cells leads to aberrant dendrite and axon morphology. (*A*) Schematic illustrating cerebellar anatomy. Purkinje cells (PC, magenta) are the only output of the cerebellar cortex. Their dendrites in the molecular layer (ML) receive inputs from granule cell parallel fibers and climbing fibers from the inferior olivary nucleus (not shown). PC cell bodies form the Purkinje cell layer (PL), while their axons project through the inner granule layer (IGL) where mature granule cells (GC) and Golgi interneurons (G) are found, to the deep cerebellar nuclei (DCN), where they synapse with the deep cerebellar nuclei neurons (DCNn). (*B-D*) *PI31^fl/fl^ PCP2-Cre* mice display progressive behavioral defects. Up to P22, *PI31^fl/fl^ PCP2-Cre* mice were not distinguishable from *PI31^fl/fl^* mice in their righting reflex and visual observation of gait. (*B*) By P30, *PI31^fl/fl^ PCP2-Cre* mice began to show loss of balance in the ledge test, which became more severe by P60. For P30, *PI31^fl/fl^* (n12) and *PI31^fl/fl^ PCP2-Cre* (n12), and for P60, *PI31^fl/fl^* (n15) and *PI31^fl/fl^ PCP2-Cre* (n24). (*C, D*) At 10 months, the gait of *PI31^fl/fl^ PCP2-Cre* mice was disturbed and mutants frequently lost balance. Gait analysis showed a significant decrease in stride length (p = 0.00258, *PI31^fl/fl^* (n4) and *PI31^fl/fl^ PCP2-Cre* (n4)). (*D*) An example of footprints used for gait analysis with measurements of length, sway and stance. Hind paw prints are blue and forepaw prints are red. The red arrow points to a common misplaced step for *PI31^fl/fl^ PCP2-Cre* mice, while the blue arrow indicates the hind paw touching the surface with splayed toes. (*E-I*) Cerebellum labeled for Calbindin (magenta) to mark Purkinje cells, Ub-GFP (green), p62 (red), and Hoechst 33342 (blue) for nuclei. Yellow box denotes inset. (*E, F*) Mature Purkinje cells at P22 appeared virtually normal in *PI31^fl/fl^ PCP2-Cre* mice when compared to *PI31^fl/fl^* mice. (*F*) PC cell bodies appeared normal and their dendritic arbors were fully developed (section of the fourth cerebellar lobe). (*G, H*) By P30, *PI31^fl/fl^ PCP2-Cre Ub-GFP* (PI31^fl/fl^ PCP2-Cre) mice Purkinje cell primary and secondary dendrites appeared swollen, while their axon terminals in the DCN showed swellings with increased Ub-GFP (yellow arrows), indicating impaired protein breakdown (*I*) Loss of PI31 caused swellings (yellow arrows) along the length of the axons in the white matter (P30 cerebellar lobe 4). (*E-H*) are single confocal slices. (*I*) is a maximum intensity projection of a confocal stack. (*E, G*) scale bars are 20 μm. (*F, H, I*) scale bars are 10 μm.

Next, we looked for evidence of changes in protein degradation and cellular structure of Purkinje cells. At P22, Purkinje cells did not appear to have morphological abnormalities in *PI31^fl/fl^ PCP2-cre* mice, with well-developed dendritic arbors and axonal morphology (Fig. 4E, F). However, we observed increased staining for Ub-GFP, a marker indicating proteotoxic stress, already at this stage (Fig. 4F). By P30, Purkinje cell primary and secondary dendrites appeared swollen in *PI31^fl/fl^ PCP2-Cre* mice (Fig. 4G). Significantly, Purkinje cell axon terminals in the DCN had swellings with increased Ub-GFP, indicating impaired protein breakdown (Fig. 4H). We also found that loss of PI31 caused swellings along the length of the axons in the white matter. These swellings were positive for Ub-GFP, indicating impaired protein breakdown in mutant PC axons (Fig. 4I). Interestingly, it appeared that the compartment most affected by the loss of PI31 at this stage was the Purkinje cell axon. By P60, the DCN contained large aggregates throughout which were positive for both poly-Ub conjugates and p62 (Fig. 5, Fig. S3). Quantification revealed a striking increase in p62-positive aggregates between 1 and 2-month old mutant mice (Fig. 5D). These data demonstrate that the loss of PI31 function exposes Purkinje cells to proteotoxic stress that progressively increases over time.

**Fig. 5:**
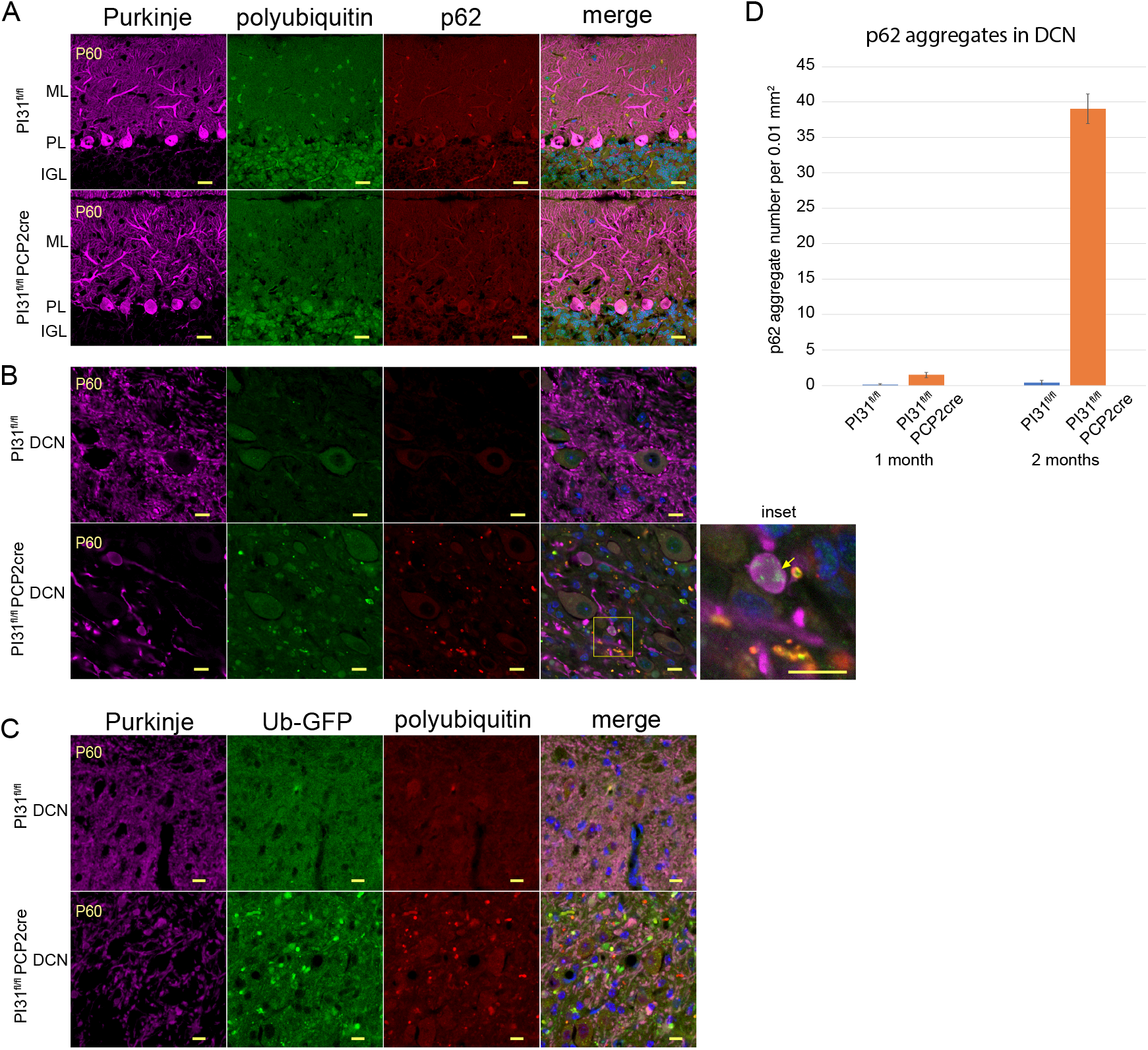
Loss of PI31 in cerebellar Purkinje cells (PCs) leads to axon degeneration and the appearance of aggregates positive for proteotoxic stress markers (p62, poly-ubiquitin and Ub-GFP). Immunostaining of cerebellum from two-month old *PI31^fl/fl^ Ub-GFP* (PI31^fl/fl^) and *PI31^fl/fl^ PCP2-Cre Ub-GFP* (PI31^fl/fl^ PCP2-Cre) mice. (*A, B*) Calbindin-staining (magenta) for Purkinje cells, poly-ubiquitin (FK2, green), p62 (red) and Hoechst 33342 (blue) for nuclei. (*A*) Loss of PI31 did not cause noticeable accumulation of either p62, FK2, or Ub-GFP aggregates in the cell bodies or dendrites of PCs. (*B*) Swellings of PC axon terminals in the DCN and accumulation of both poly-Ub and p62-positive aggregates was observed in the DCN of P60 mutant mice (see also Supp. Figure 3). Yellow arrow in inset points to an example of poly-Ub aggregates (green) in axon swellings. (*C*) Ub-GFP-positive aggregates (green) in the DCN of P60 mutants. Images in (*A-C*) are single confocal slices. (*A*) scale bars are 20 μm. (*B,C*) scale bars are 10 μm. (*D*) Quantification of p62 aggregates in the DCN reveals a dramatic increase from one to two-month old *PI31^fl/fl^ PCP2-Cre mice*. p62-positive aggregates were counted in a 0.01 mm^2^ area in the DCN of P30 *PI31^fl/fl^* (n3) and *PI31^fl/fl^ PCP2-Cre* (n3) and P60 *PI31^fl/fl^* (n4) and *PI31^fl/fl^ PCP2-Cre* (n4) mice. Statistical significance between loss of PI31 mice and control mice at 2 months is p=6.49 x 10^-5^ (two-tailed T test).

Many neurodegenerative diseases are associated with gliosis (glial cell reactivation and proliferation), which often precedes the formation of aggregates, tangles and plaques (71, 79). An increase in glial fibrillary acid-protein (GFAP) and hypertrophy of astrocytes are markers of astrogliosis (80). In the mouse cerebellum, GFAP mainly stains the Bergmann glia in the Purkinje and molecular layers, and astrocytes which are predominantly found in the white matter. For *PI31^fl/fl^ PCP2-cre* mice, the GFAP staining at P22 was very similar to their control littermates (Fig. S4). However, by P30, there was a dramatic increase in reactive astrocytes in the DCN, but not the Purkinje or molecular layers (Fig. S4). This indicates that the Purkinje cell axon terminals are damaged in such a way as to induce gliosis at this time. At P60, we observed some reactive astrocytes in the Purkinje layer, coinciding with the onset of Purkinje cell loss.

An extension of this analysis to older mice revealed that mutant Purkinje cells had lost almost all their axon terminals in the DCN by the age of 10 months (Fig. 6A). We also observed axonal torpedoes, focal swellings on Purkinje cell axons that have long been associated with cerebellar disorders (Fig. 6B) (81–84). In addition, at this time there was a major reduction in Purkinje cell number (Fig. 6B, C). This striking loss of Purkinje cells upon inactivation of *PI31* is consistent with the motor defects revealed by gait analysis in these mice. Collectively, our results indicate that loss of PI31 initially impairs local protein homeostasis in axons and dendrites of Purkinje cells and disrupts their structure, and that over time this compromises cell survival.

**Fig. 6:**
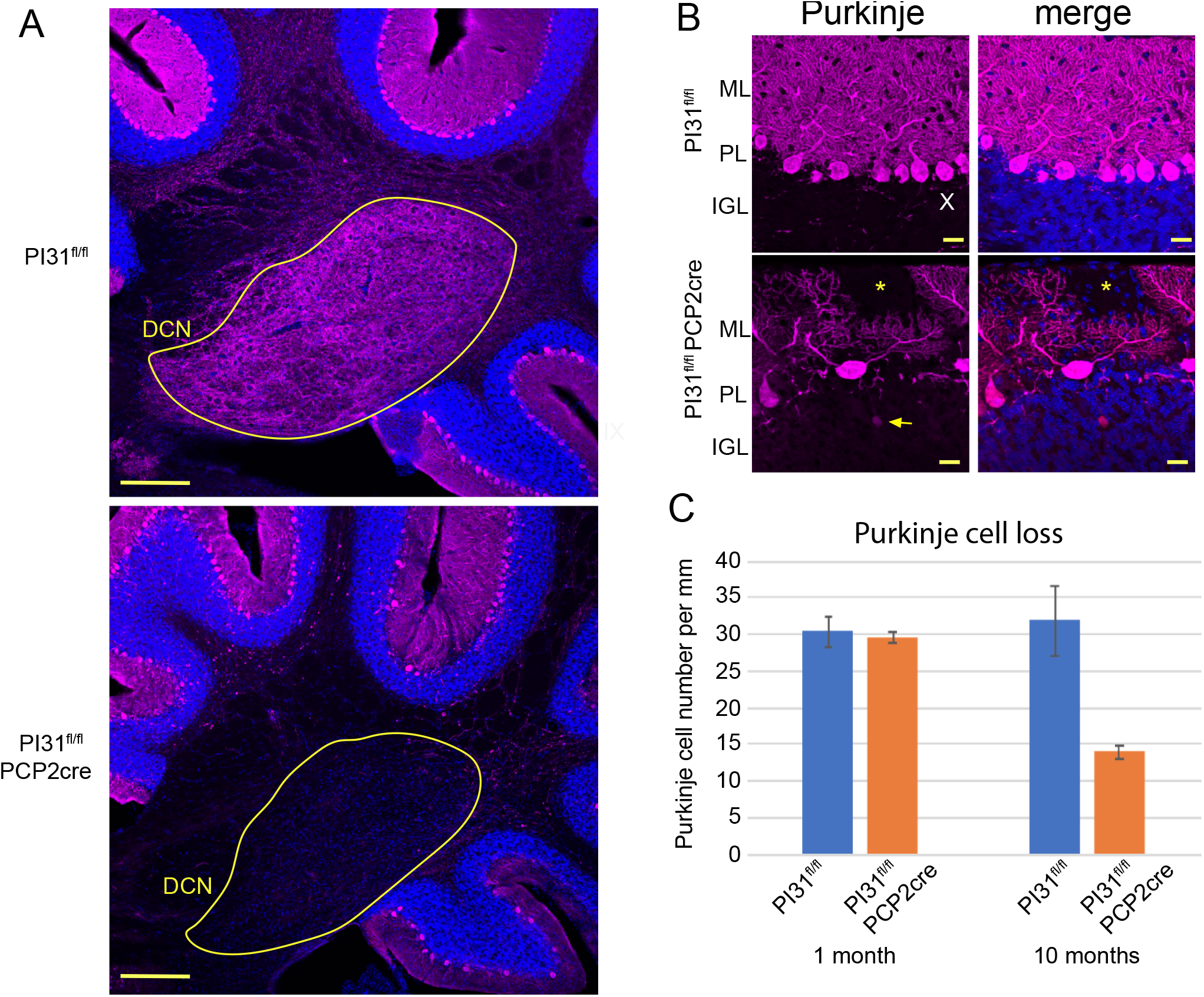
Loss of PI31 in cerebellar Purkinje cells leads to Purkinje cell loss. (*A*) Overview showing loss of Purkinje cell axons in the DCN of the *PI31^fl/fl^ PCP2-Cre* cerebellum. Immunostaining of cerebellum from 10 month old *PI31^fl/fl^* and *PI31^fl/fl^ PCP2-Cre* mice. PCs were marked by Calbindin staining (magenta) and nuclei with Hoechst 33342 (blue). Scale bar 200 μm. DCN circled in yellow. Note near complete loss of Calbindin staining in the DCN of the *PI31^fl/fl^ PCP2-Cre* cerebellum. (*B*) Aberrant morphology and loss of Purkinje cells. The yellow asterisk marks a gap of PC dendrites in the molecular layer, and the arrow points to an axonal torpedo. PC dendritic arbors were also severely reduced. PCs were marked by Calbindin (magenta) and nuclei with Hoechst 33342 (blue). Maximum intensity projection (6 μm). Scale bar 20 μm. (*C*) Graph showing number of PCs per mm of Purkinje layer. At 1 month of age the number of PCs was similar to control, but by 10 months PC number had decreased to about 44% of control mice.

## Discussion

In the present study, we addressed the physiological role of PI31 (“<u>p</u>roteasome inhibitor of <u>31</u>kD”) in mammals by generating complete and conditional PI31 mouse mutants. In contrast to the function implied by its original name, we provide compelling evidence for a role of PI31 in stimulating protein breakdown that is critical for maintaining synaptic structures and long-term survival of neurons. Interestingly, the first morphological defects caused by conditional inactivation of PI31, in both motor neurons and Purkinje cells of the cerebellum, were observed in the distal part of neurons, namely axons and synapses. These structural defects were accompanied by molecular proteotoxic stress markers (Ub-GFP, FK2 and P62) at the NMJ and in deep cerebellar nuclei (DCN). Importantly, symptoms became progressively more severe with age, culminating in the degeneration of axons and dendrites, and eventually cell loss. These morphological abnormalities were also associated with motoric problems, modeling ALS and cerebellar disorders, respectively. Overall, these observations suggest that PI31 activity is required for protein homeostasis in distal compartments of neurons.

A plausible mechanistic explanation for these defects comes from our recent finding that PI31 serves as an adapter protein to mediate fast axonal transport of proteasomes (43). PI31 can directly bind to dynein light chain proteins and thereby couples proteasomes to cellular motors. Significantly, inactivation of PI31 in *Drosophila* blocks proteasome motility and disrupts synaptic protein homeostasis and structure. This function of PI31 is conserved from insects to mammals, since inactivation of PI31 in mouse hippocampal neurons impairs proteasome movement in axons as well (43). Although we have not yet been able to perform live cell imaging analyses to directly demonstrate a requirement of PI31 for proteasome movement in mouse motor neurons or Purkinje cells, impaired transport of proteasomes in axons and dendrites provides an attractive explanation for the observed defects in protein homeostasis at nerve endings. Synapses are highly dynamic sites of active local protein synthesis, which has to be balanced by local protein degradation (85). According to our model, inactivation of PI31 leads to a shortage of proteasomes at synapses, which in turn causes the accumulation of poly-Ub proteins and p62-positive aggregates. It appears that backup clearance mechanisms, such as autophagy and possibly also shedding of microvesicles/exosomes, can partially compensate for the loss of PI31-mediated proteasome activity in PI31 mutants (6, 43, 86–90). The latter would also explain the gliosis that we observed already at P30 in *PI31^fl/fl^ PCP2-Cre* mice. However, even if backup clearance pathways are induced, neurons eventually succumb to defects in the absence of PI31 function.

Interestingly, conditional inactivation of PI31 in motor neurons resulted in kyphosis and ALS-like behavioral phenotypes comparable to inactivation of Rpt3, an essential component of the proteasome 19S regulatory particle (54). This is somewhat surprising, since Rpt3, unlike PI31, is an essential component of the proteasome regulatory particle. In the mouse embryo, complete inactivation of Rpt3 causes a much more severe phenotype (*Rpt3-deficient* mice die before implantation) than complete inactivation of PI31 (69). Unlike PI31, inactivation of Rpt3 primarily disrupted protein homeostasis in the motor neuron cell body (54). We interpret this to indicate the importance of localized proteasome activity in axons and dendrites of neurons: whereas Rpt3 function is required in every cellular compartment, including the nucleus and cell body, PI31-mediated proteasome transport appears to be particularly important for protein homeostasis in axons and dendrites. Since neuronal activity critically relies on proper synaptic function, it may be particularly sensitive to perturbation of synaptic protein homeostasis mediated by PI31. Moreover, it is also possible that proteasomes can be active independent of the main regulatory particle, at least to a certain degree and in some subcellular compartments. Together, the critical and conserved role of PI31 in proteasome transport, the accumulation of non-degraded poly-Ub conjugates and induction of proteotoxic stress in PI31 mutants, and comparison with Rpt3 inactivation in motor neurons, all support a function of PI31 as a positive regulator of proteasome activity.

The activity of PI31 critically depends on Fbxo7, also termed PARK15 in humans, which encodes the substrate-recognition component of an SCF E3-ligase complex and binds directly to PI31 (26, 30–32). Conditional inactivation of PI31 in mice causes neuronal degeneration, and human mutations in Fbxo7/PARK15 are associated with neurodegenerative diseases (33, 34, 42, 91, 92). Strikingly, inactivation of Fbxo7 leads to severe reduction of PI31 protein and reduced proteasome activity in both *Drosophila* and mice (26, 32, 33). This suggests the possibility of a direct mechanistic link between PI31, reduced proteasome function and human disease. However, a role for PI31 in mediating the proteasome impairment and neuronal degeneration observed in Fbxo7/PARK15 mutants was dismissed based on results from cell culture experiments (33, 93). The current study, together with the work of Liu et al. (2019), suggest that this topic deserves more careful examination (43). MEFs deficient for PI31 survived for extended periods of time, and we were unable to detect a major reduction of total proteasome activity in these cells. However, local defects in proteasome activity would not have been revealed by these approaches. When combined with proteasome inhibitors, inactivation of PI31 resulted in a small but significant reduction of cell survival. These findings are consistent with the reported role of the yeast PI31 ortholog to alleviate proteotoxic stress (27). We also saw that PI31-Null MEFs had slight defects in proteasome assembly, as indicated by reduced numbers of single-capped 26S particles. Therefore, PI31 may regulate proteasomes by multiple mechanisms: through the assembly of functional 26S proteasomes from 19S and 20S particles, and via transport (29, 43). In any event, it appears that neurons are particularly sensitive to the loss of PI31 function, and a neuro-protective role of PI31 should not be ruled out based on results with non-neuronal immortalized cells.

A neuro-protective role of PI31 was particularly obvious in motor neurons and Purkinje cells, two major types of neurons with long axons. Therefore, it is possible that large neurons axons are especially sensitive to the impairment of PI31-mediated proteasome transport since they face a particularly challenging problem in allocating proteasomes to compartments where protein breakdown has to occur. Moreover, Purkinje cells are very sensitive to proteotoxic stress, and Purkinje cell pathology is a common finding in cerebellar diseases (47, 71, 79). Consistent with this, inactivation of PI31 in these cells caused proteotoxic stress as soon as these cells matured (P22), and it led to behavioral and anatomical defects that are characteristic for cerebellar diseases. These include progressively more severe loss of balance and defects in gait, which coincided with axonal swelling, torpedoes, gliosis and eventually loss of Purkinje cell bodies.

It is well known that protein clearance mechanisms decline during the aging process, and impairment of the UPS in neurodegenerative diseases has been widely reported (35, 36, 39, 40, 90, 94–97). Most of the attention in previous studies has been focused on the ability of aggregates to inhibit proteasome activity (37, 41). However, aggregate-mediated proteasome inhibition cannot explain the failure to degrade poly-Ub proteins and accumulate aggregates in the first place. We suggest that impaired activity of PI31, caused either by mutations affecting PI31 function (such as Fbxo7/PARK15), or through epigenetic mechanisms during the normal aging process, contributes to age-related neuronal degeneration. The mouse mutant strains described here provide powerful models to further investigate this possibility.

## Acknowledgement

We thank Drs. David Ng and Thomas Jessell for advice, invaluable support and for providing the Hb9-Cre strain. We are also grateful to Drs. Thomas Misgeld and Monika Leischner for their helpful advice regarding the analysis of muscle innervation. We thank members of the Steller and Hatten labs for thoughtful comments and helpful advice throughout. This work was supported by NIH grant RO1GM60124 and a grant from the Cure Alzheimer’s Foundation to H.S, The Rockefeller University Center for Clinical and Translational Science to A.L., and support from the Eugene W. Chinery 2012 Trust to E.E.G. and M.E.H.

## Author Contributions

Conceptualization, A.M., J.R., A.L., K.L. and H.S.; Investigation and analysis, A.M., J.R., A.L., K.L., E.E.G., M.E.H. and H.S. Resources, H.S.; Writing, A.M., J.R., E.E.G., M.E.H., and H.S.; Funding Acquisition, H.S.; Supervision, H.S.

## Materials and Methods

### Mice

In order to generate PI31 mutant using CRISPR\Cas9 technology we have designed a single-guide targeting RNA that will target the mouse PSMF1 gene in the first exon right after the first Methionine (Psmf1a-1 CACCGAACGGCTACTATGCCTT, Psmf1a-2 AAACAAGGCATAGTAGCCGTTC). Targeting sequence was ordered as oligos and cloned into plasmid px330 (A gift from Dr. Zhang lab) that contains both Cas9 and the tracer RNA sequence. ES cell were transfected with this plasmid and Surveyor assay was performed to assess targeting efficiency. Gene targeting resulted in 5 mutant lines, all of which were heterozygous for an indel in the PSMF1 gene. After sequencing the relevant genomic loci we have decided to continue our research with one clone, which have a 16bp deletion in the first exon (nt103-118 of ORF) that results in a frame shift and a premature stop codon in the second exon (nt239). Homozygous mutants are not viable and die perinatal; hence, we used tissue from E13 homozygous mutant embryos (which are morphologically similar to wild types) to test the presence of PI31 protein.

For the generation of conditional PI31 knockout mice, we have purchased commercially available ES cell (Psmf1^tm1a(EUCOMM)Hmgu^). We sequenced these ES cells to validate they harbor PI31 knockout first allele with conditional potential. Chimeras were generated by injection of targeted ES cells into albino blastocysts purchased from The Jackson Laboratory (B6(Cg)-Tyr<c-2J>/J). Next chimeras were crossed to B6 albino mice (B6(Cg)-Tyrc-2J/J) from The Jackson Laboratory. Heterozygous transgene mice were then bred to mouse expressing FLP-1 recombinase gene under the direction of the human ACTB promoter (The Jackson Laboratory #005703), to create a mouse line where the third exon PI31 is flanked by two LoxP sites (PI31^fl/fl^). Conditional PI31^fl/fl^ were generated using CDX2-Cre, Hb9-Cre, Ubc-CreERt, purchased from The Jackson Laboratory. C57B1/6J *PI31^fl/fl^* mice were crossed to C57B1/6J *Tg(Pcp2-Cre)GN135Gsat* generated by and obtained from GENSAT (Rockefeller University). *PI31^fl/fl^Pcp2-Cre* mice were compared to *PI31^fl/fl^* control littermates. Genotyping of *PI31^fl/fl^* and *Pcp2-Cre* alleles was carried out by PCR. All animal work was performed as required by the United States Animal Welfare Act and the National Institutes of Health’s policy to ensure proper care and use of laboratory animals for research, and under established guidelines and supervision by the Institutional Animal Care and Use Committee (IACUC) of The Rockefeller University. Mice were housed in accredited facilities of the Association for Assessment of Laboratory Animal Care (AALAC) in accordance with the National Institutes of Health guidelines.

### Behavioral Tests

#### Ledge test

The mouse was placed on the ledge of the cage and scored for coordination and balance (98). A mouse which walked the ledge with few slips and descended into the cage gracefully, landing on its paws, was given a score of 0. A mouse which lost its footing received a score of 1. A mouse that did not use its hind paws well and landed on its head instead of its paws when descending into the cage received a score of 2. A mouse that refused to move, even with gentle encouragement, or fell or nearly fell while walking the ledge or lowering itself into the cage received a score of 3.

#### Gait Analysis

The fore paws of each mice were painted with nontoxic red paint and the hind paws were painted with nontoxic blue paint, after which the mice were allowed to walk on paper along a 10 cm wide by 76 cm long corridor with 20 cm high opaque walls. This was done three times for each mouse and each parameter was measured for the three longest strides in the middle of a run. The footprint patterns were analyzed for three step parameters (all measured in centimeters). As per Carter et al., four parameters were measured: (1) Stride length was measured as the average distance of forward movement between each stride. (2) Hind-base width, or sway, was measured as the average distance between left and right hind footprints. These values were determined by measuring the perpendicular distance of a given step to a line connecting its opposite preceding and proceeding steps. (3) Distance from left hind footprint to right hind was measured to determine stance length (99).

### Native gel electrophoresis

Protein extracts of tissue were prepared in buffer (50 mM Tris-HCl (pH 8.0), 5 mM MgCl2, 0.5 mM EDTA, 2 mM ATP, 0.2% NP4, protease inhibitor, phosphatase inhibitor (Pfizer), using liquid nitrogen freezing and thawing technique. Extracts were cleared by centrifugation at 14, 000 RPM for 30 min to remove nuclei and cell debris, and protein concentrations were measured by Bradford assay. To resolve proteasomes, 26-well 3–8% Tri-acetate gels (Criterion) were used. Samples were mixed with 2X native loading buffer (Bio-Rad) just before loading. Electrophoresis were carried out at RT (1 hour at 50V) and then at 4°C for additional 5 hours at 120V, in running buffer (0.45M Tris, 0.45M Boric Acid, 5mM EDTA, 12.5 mM MgCl2, supplied with 0.5 mM DTT, 0.5 mM ATP). For immunoblotting, proteins in native gels were transferred to 0.45 mm PVDF membranes. 26S and 20S were detected with anti-Alpha7 antibodies (1:5000, Enzo). Purified bovine 20S, and 26S proteasomes (UBPBio), were used as standards.

### Antibodies and Western blot

Protein extracts were prepared as for Native gel electrophoresis. For western blotting, proteins were resolved by SDS-PAGE and transferred onto a 0.45 mm PVDF membrane. Membranes were blocked overnight at 4°C with 5% milk in phosphate buffered saline (PBS) and 0.5% Tween-20 (PBST). Membranes were incubated for 60 min with one of the following antibodies: FK2 (Enzo, 1:2000), PI31 (Thermo Fisher, 1:2000), Actin HRP conjugated (Cell Signaling,1:10000), Tubulin. Primary antibodies were detected by secondary species-specific HRP-congugated antibodies (Jackson ImmunoResearch, 1:5000). Detection was performed with Amersham ECL Western Blotting Detection Reagent (GE Healthcare).

### Cell Survival Assay

Low cycle (4–6) primary mouse embryonic fibroblasts from wild type or PI31 KO embryos were seeded in a 96 well plate at a concentration of 15,000 cells per well and incubated over night in DMEM containing 10% FBS and penicillin/streptomycin at 37°C under 5% CO2 (v/v). Then, cells were incubated for additional 18-22 hours with different concentration of the proteasome inhibitors Bortezomib or MG132, three wells per concentration. Cell viability was measured using PrestoBlue viability reagent (molecular probed, life technologies) as described in the manufacture protocol. Experiment was repeated three independent times.

### Immunofluorescence Muscle innervation

Whole mount staining of triangularis sterni muscle was done as described in Brill et al., Neuron 2016 (100). Briefly, the thoracic wall containing the triangularis sterni muscle was dissected and fixed in 4% PFA for 1 hour on ice. The fixed tissue was then washed for 10 minuets in 0.1M glycine and the triangularis sterni muscle dissected out. The dissected muscle was transferred into blocking solution (10% goat serum, 0.5% TX-100 in PBS) for 1 hour at room temperature. After blocking, tissue was incubated with Alexa488 conjugated primary antibodies for Tuj1 (mouse monoclonal, 1:200; BD Pharmingen #560339) and Synapsin (Rabbit monoclonal, 1:200; CST #13197) over night at 4c. To lable post-synaptic acetylcholine receptors, samples were incubated in Alexa594-alpha-bungarotoxin (Invitrogen, B-13423, 50μg/ml, 1:500 in PBS). Tissue was washed 3x in PBS and mounted on a slide with Vectashield mounting media. <u>Quantification of axonal swellings</u> - for the quantification of axonal swellings, z-stack images of age matched PI31^f/f^ controls and PI31^f/f^ Hb9:Cre KO animals were taken at low magnification (10x). Maximal intensity Z-projects images were generated using Fiji software and axonal swellings were counted manually. Average number of axonal swelling was calculated based on 3 images per animal. Image size – 850□m^2^. 1MO – control n= 3, KO n=3, 3MO – control n=2, KO n=3, 5MO – control n=2, KO n=4. P-Value was calculated using the student t-test.

### Immunofluorescence of cerebellar sections

Mice were perfused with 4% PFA/PBS, the brains were removed and then postfixed in 4% PFA/PBS overnight. They were then washed in PBS, soaked in 15% sucrose/PBS for several hours until they sank, and then soaked in 30% sucrose/PBS overnight. The brains were then embedded in Richard-Allan Scientific Neg-50^™^ Frozen Section Medium (Thermo Scientific) and 50 μm sagittal sections were made using a Leica CM 3050S cryostat.

Primary antibodies used for immunofluorescence included rabbit anti-GFP (Invitrogen, 1/2000), mouse anti-Calbindin (CALB1) (Swant, 1/1000), rabbit anti-Calbindin, (1/1000), rabbit anti-NeuN (Abcam), mouse anti-polyubiquitin (FK2) (Enzo, 1/200) and mouse anti-SQSTM1/p62 (Abcam, 1/500). Secondary antibodies used included the species appropriate Alexa Fluor^®^ 488, Alexa Fluor^®^ 568 and Alexa Fluor^®^ 647 (Invitrogen, 1/500).

The floating cryostat sections were permeabilized and blocked overnight in 1x PBS/0.3% Triton X-100/5% Normal Donkey Serum (NDS). The sections were then incubated overnight with primary antibodies diluted in 1x PBS/0.3% Triton X100/5% NDS. Subsequently, the sections were washed 3 times for 20 minutes each in 1x PBS/0.3% Triton X-100/5% NDS and incubated overnight with Alexa Fluor^®^ conjugated secondary antibodies diluted in 1x PBS/0.3% Triton X-100/5% NDS. The sections were then washed 4 times for 30 minutes in 1x PBS/0.3% Triton X-100/5% NDS and then mounted with Molecular Probes ProLong^®^ Diamond anti-fade mounting medium (Invitrogen). Images where captured on a Zeiss confocal LSM780 with Zen software. Images shown are single slices unless otherwise noted.

### Purkinje cell counting

Purkinje cell bodies were counted from confocal images of cerebellar sections immunostained with anti-CALB1 to mark the Purkinje cells of 1 and 10 month old *PI31^fl/fl^ Pcp2-Cre* mice (n3) and their control littermates (n3). This count was converted to cells/mm^2^.

### P62 Aggregates

Confocal images of cerebellar sections were stained with anti-CALB1 to mark the Purkinje cells and anti-p62. Aggregates within the DCN were counted and converted to aggregates/area for P30 and P60 *PI31^fl/fl^ Pcp2-Cre* mice (n3) and their control littermates (n3).

## REFERENCES

1. A. Hershko, A. Ciechanover, The ubiquitin system. Annual review of biochemistry 67, 425–479 (1998).

2. M. H. Glickman, A. Ciechanover, The ubiquitin-proteasome proteolytic pathway: destruction for the sake of construction. Physiological reviews 82, 373–428 (2002).

3. A. L. Goldberg, On prions, proteasomes, and mad cows. The New England journal of medicine 357, 1150–1152 (2007).

4. I. Dikic, Z. Elazar, Mechanism and medical implications of mammalian autophagy. Nat Rev Mol Cell Biol 19, 349–364 (2018).

5. J. Labbadia, R. I. Morimoto, The biology of proteostasis in aging and disease. Annu Rev Biochem 84, 435–464 (2015).

6. S. Wolff, J. S. Weissman, A. Dillin, Differential scales of protein quality control. Cell 157, 52–64 (2014).

7. A. Varshavsky, The ubiquitin system, an immense realm. Annu Rev Biochem 81, 167–176 (2012).

8. B. Levine, G. Kroemer, Autophagy in the pathogenesis of disease. Cell 132, 27–42 (2008).

9. W. Baumeister, J. Walz, F. Zuhl, E. Seemuller, The proteasome: paradigm of a self-compartmentalizing protease. Cell 92, 367–380 (1998).

10. G. A. Collins, A. L. Goldberg, The Logic of the 26S Proteasome. Cell 169, 792–806 (2017).

11. A. S. Hafner, P. G. Donlin-Asp, B. Leitch, E. Herzog, E. M. Schuman, Local protein synthesis is a ubiquitous feature of neuronal pre- and postsynaptic compartments. Science 364 (2019).

12. A. Biever, P. G. Donlin-Asp, E. M. Schuman, Local translation in neuronal processes. Curr Opin Neurobiol 57, 141–148 (2019).

13. B. Bingol, M. Sheng, Deconstruction for reconstruction: the role of proteolysis in neural plasticity and disease. Neuron 69, 22–32 (2011).

14. D. S. Campbell, C. E. Holt, Chemotropic responses of retinal growth cones mediated by rapid local protein synthesis and degradation. Neuron 32, 1013–1026 (2001).

15. A. N. Hegde, K. A. Haynes, S. V. Bach, B. C. Beckelman, Local ubiquitin-proteasome-mediated proteolysis and long-term synaptic plasticity. Front Mol Neurosci 7, 96 (2014).

16. C. T. Kuo, L. Y. Jan, Y. N. Jan, Dendrite-specific remodeling of Drosophila sensory neurons requires matrix metalloproteases, ubiquitin-proteasome, and ecdysone signaling. Proc Natl Acad Sci U S A 102,15230–15235 (2005).

17. R. J. Watts, E. D. Hoopfer, L. Luo, Axon pruning during Drosophila metamorphosis: evidence for local degeneration and requirement of the ubiquitin-proteasome system. Neuron 38, 871–885 (2003).

18. J. J. Yi, M. D. Ehlers, Ubiquitin and protein turnover in synapse function. Neuron 47, 629–632 (2005).

19. H. C. Tai, E. M. Schuman, Ubiquitin, the proteasome and protein degradation in neuronal function and dysfunction. Nat Rev Neurosci 9, 826–838 (2008).

20. C. T. Kuo, S. Zhu, S. Younger, L. Y. Jan, Y. N. Jan, Identification of E2/E3 ubiquitinating enzymes and caspase activity regulating Drosophila sensory neuron dendrite pruning. Neuron 51, 283–290 (2006).

21. M. Schmidt, J. Hanna, S. Elsasser, D. Finley, Proteasome-associated proteins: regulation of a proteolytic machine. Biol Chem 386, 725–737 (2005).

22. H. C. Besche, A. Peth, A. L. Goldberg, Getting to first base in proteasome assembly. Cell 138, 25–28 (2009).

23. A. Rousseau, A. Bertolotti, Regulation of proteasome assembly and activity in health and disease. Nat Rev Mol Cell Biol 19, 697–712 (2018).

24. M. Chu-Ping, C. A. Slaughter, G. N. DeMartino, Purification and characterization of a protein inhibitor of the 20S proteasome (macropain). Biochimica et biophysica acta 1119, 303–311 (1992).

25. S. L. McCutchen-Maloney et al., cDNA cloning, expression, and functional characterization of PI31, a proline-rich inhibitor of the proteasome. The Journal of biological chemistry 275,18557–18565 (2000).

26. M. Bader et al., A conserved F box regulatory complex controls proteasome activity in Drosophila. Cell 145, 371–382 (2011).

27. H. Yashiroda et al., N-Terminal alpha7 Deletion of the Proteasome 20S Core Particle Substitutes for Yeast PI31 Function. Molecular and cellular biology 35, 141–152 (2015).

28. B. J. Yang et al., Arabidopsis PROTEASOME REGULATOR1 is required for auxin-mediated suppression of proteasome activity and regulates auxin signalling. NatCommun 7, 11388 (2016).

29. P. F. Cho-Park, H. Steller, Proteasome regulation by ADP-ribosylation. Cell 153, 614–627(2013).

30. R. Kirk et al, Structure of a conserved dimerization domain within the F-box protein Fbxo7 and the PI31 proteasome inhibitor. The Journal of biological chemistiy 283, 22325–22335 (2008).

31. D. E. Nelson, S. J. Randle, H. Laman, Beyond ubiquitination: the atypical functions of Fbxo7 and other F-box proteins. Open Biol 3, 130131 (2013).

32. M. Bader, E. Arama, H. Steller, A novel F-box protein is required for caspase activation during cellular remodeling in Drosophila. Development 137, 1679–1688 (2010).

33. S. Vingill et al., Loss of FBXO7 (PARK15) results in reduced proteasome activity and models a parkinsonism-like phenotype in mice. EMBO J 35, 2008–2025 (2016).

34. S. R. W. Stott et al., Loss of FBXO7 results in a Parkinson’s-like dopaminergic degeneration via an RPL23-MDM2-TP53 pathway. J Pathol 10.1002/path.5312 (2019).

35. A. Ciechanover, P. Brundin, The ubiquitin proteasome system in neurodegenerative diseases: sometimes the chicken, sometimes the egg. Neuron 40, 427–446 (2003).

36. N. P. Dantuma, L. C. Bott, The ubiquitin-proteasome system in neurodegenerative diseases: precipitating factor, yet part of the solution. Front Mol Neurosci 7, 70 (2014).

37. N. Myeku et al., Tau-driven 26S proteasome impairment and cognitive dysfunction can be prevented early in disease by activating cAMP-PKA signaling. Nat Med 22, 46–53 (2016).

38. J. N. Keller, K. B. Hanni, W. R. Markesbery, Impaired proteasome function in Alzheimer’s disease. J Neurochem 75, 436–439 (2000).

39. D. C. Rubinsztein, The roles of intracellular protein-degradation pathways in neurodegeneration. Nature 443, 780–786 (2006).

40. M. Schmidt, D. Finley, Regulation of proteasome activity in health and disease. Biochim Biophys Acta 1843, 13–25 (2014).

41. T. A. Thibaudeau, R. T. Anderson, D. M. Smith, A common mechanism of proteasome impairment by neurodegenerative disease-associated oligomers. NatCommun 9, 1097 (2018).

42. S. Conedera et al., FBXO7 mutations in Parkinson’s disease and multiple system atrophy. Neurobiol Aging 40, 192 el91–192 el95 (2016).

43. K. Liu et al., PI31 is an adaptor protein for proteasome transport in axons and required for synaptic development. Dev Cell 10.1016/j.devcel.2019.06.009 (2019).

44. K. C. Kanning, A. Kaplan, C. E. Henderson, Motor neuron diversity in development and disease. Annu Rev Neurosci 33, 409–440 (2010).

45. M. E. Hatten, N. Heintz, Mechanisms of neural patterning and specification in the developing cerebellum. Annu Rev Neurosci 18, 385–408 (1995).

46. S. L. Palay, V. Chan-Palay, Cerebellar cortex: cytology and organization (Springer, Berlin, Heidelberg, New York, 1974], pp. xii, 348 p.

47. M. Huang, D. S. Verbeek, Why do so many genetic insults lead to Purkinje Cell degeneration and spinocerebellar ataxia? Neurosci Lett 688, 49–57 (2019).

48. C. Rothblum-Oviatt et al., Ataxia telangiectasia: a review. Orphanet J Rare Dis 11, 159 (2016).

49. E. D. Louis, P. L. Faust, J. P. Vonsattel, Purkinje cell loss is a characteristic of essential tremor. Parkinsonism Relat Disord 17, 406–409 (2011).

50. S. H. Fatemi et al., Consensus paper: pathological role of the cerebellum in autism. Cerebellum 11, 777–807 (2012).

51. J. Nijssen, L. H. Comley, E. Hedlund, Motor neuron vulnerability and resistance in amyotrophic lateral sclerosis. Acta Neuropathol 133, 863–885 (2017).

52. L. Ferraiuolo, J. Kirby, A. J. Grierson, M. Sendtner, P. J. Shaw, Molecular pathways of motor neuron injury in amyotrophic lateral sclerosis. Nat Rev Neurol 7, 616–630 (2011).

53. J. P. Taylor, R. H. Brown, Jr., D. W. Cleveland, Decoding ALS: from genes to mechanism. Nature 539, 197–206 (2016).

54. Y. Tashiro et al., Motor neuron-specific disruption of proteasomes, but not autophagy, replicates amyotrophic lateral sclerosis. The Journal of biological chemistiy 287, 42984–42994 (2012).

55. M. Ding, C. Weng, S. Fan, Q. Cao, Z. Lu, Purkinje Cell Degeneration and Motor Coordination Deficits in a New Mouse Model of Autosomal Recessive Spastic Ataxia of Charlevoix-Saguenay. Front Mol Neurosci 10, 121 (2017).

56. T. Unno et al., Development of Purkinje cell degeneration in a knockin mouse model reveals lysosomal involvement in the pathogenesis of SCA6. Proceedings of the National Academy of Sciences of the United States of America 109, 17693–17698 (2012).

57. J. Cendelin, From mice to men: lessons from mutant ataxic mice. Cerebellum Ataxias 1, 4 (2014).

58. L. Ljungberg et al., Transient Developmental Purkinje Cell Axonal Torpedoes in Healthy and Ataxic Mouse Cerebellum. Front Cell Neurosci 10, 248 (2016).

59. G. Testa et al., A reliable lacZ expression reporter cassette for multipurpose, knockout-first alleles. Genesis 38, 151–158 (2004).

60. W. C. Skarnes et al., A conditional knockout resource for the genome-wide study of mouse gene function. Nature 474, 337–342 (2011).

61. M. Fujimuro, H. Sawada, H. Yokosawa, Dynamics of ubiquitin conjugation during heat-shock response revealed by using a monoclonal antibody specific to multi-ubiquitin chains. European journal of biochemistry / FEBS 249, 427–433 (1997).

62. Y. Katsuragi, Y. Ichimura, M. Komatsu, p62/SQSTM1 functions as a signaling hub and an autophagy adaptor. FEBS J 282, 4672–4678 (2015).

63. J. Moscat, M. T. Diaz-Meco, p62 at the crossroads of autophagy, apoptosis, and cancer. Cell 137,1001–1004 (2009).

64. A. L. Goldberg, Development of proteasome inhibitors as research tools and cancer drugs. The Journal of cell biology 199, 583–588 (2012).

65. S. Arber et al., Requirement for the homeobox gene Hb9 in the consolidation of motor neuron identity. Neuron 23, 659–674 (1999).

66. X. Yang et al., Patterning of muscle acetylcholine receptor gene expression in the absence of motor innervation. Neuron 30, 399–410 (2001).

67. D. M. Anderson et al., Severe muscle wasting and denervation in mice lacking the RNA-binding protein ZFP106. Proceedings of the National Academy of Sciences of the United States of America 113, E4494–4503 (2016).

68. P. McGoldrick, P. I. Joyce, E. M. Fisher, L. Greensmith, Rodent models of amyotrophic lateral sclerosis. Biochimica et biophysica acta 1832, 1421–1436 (2013).

69. Y. Sakao et al., Mouse proteasomal ATPases Psmc3 and Psmc4: genomic organization and gene targeting. Genomics 67, 1–7 (2000).

70. C. J. Cummings et al., Over-expression of inducible HSP70 chaperone suppresses neuropathology and improves motor function in SCA1 mice. Hum Mol Genet 10, 1511–1518 (2001).

71. S. M. Ronnebaum, C. Patterson, J. C. Schisler, Emerging evidence of coding mutations in the ubiquitin-proteasome system associated with cerebellar ataxias. Hum Genome Var 1, 14018 (2014).

72. K. Sasaki et al., PAC1 gene knockout reveals an essential role of chaperone-mediated 20S proteasome biogenesis and latent 20S proteasomes in cellular homeostasis. Mol Cell Biol 30, 3864–3874 (2010).

73. M. Komatsu et al., Essential role for autophagy protein Atg7 in the maintenance of axonal homeostasis and the prevention of axonal degeneration. Proc Natl Acad Sci U S A 104, 14489–14494 (2007).

74. J. Nishiyama, E. Miura, N. Mizushima, M. Watanabe, M. Yuzaki, Aberrant membranes and double-membrane structures accumulate in the axons of Atg5-null Purkinje cells before neuronal death. Autophagy 3, 591–596 (2007).

75. C. Wang, C. C. Liang, Z. C. Bian, Y. Zhu, J. L. Guan, FIP200 is required for maintenance and differentiation of postnatal neural stem cells. Nat Neurosci 16, 532–542 (2013).

76. V. Redmann et al., Clec16a is Critical for Autolysosome Function and Purkinje Cell Survival. Sci Rep 6, 23326 (2016).

77. M. Kim et al., Mutation in ATG5 reduces autophagy and leads to ataxia with developmental delay. Elife 5 (2016).

78. D. H. Margolin et al., Ataxia, dementia, and hypogonadotropism caused by disordered ubiquitination. N Engl J Med 368,1992–2003 (2013).

79. J. Vriend, S. Ghavami, H. Marzban, The role of the ubiquitin proteasome system in cerebellar development and medulloblastoma. Mol Brain 8, 64 (2015).

80. M. V. Sofroniew, Astrogliosis. Cold Spring Harb Perspect Biol 7, a020420 (2014).

81. E. D. Louis et al., Neuropathologic findings in essential tremor. Neurology 66, 1756–1759 (2006).

82. Q. Yang et al., Morphological Purkinje cell changes in spinocerebellar ataxia type 6. Acta neuropathologica 100, 371–376 (2000).

83. E. D. Louis, S. H. Kuo, J. P. Vonsattel, P. L. Faust, Torpedo formation and Purkinje cell loss: modeling their relationship in cerebellar disease. Cerebellum 13, 433–439 (2014).

84. A. Hirano, H. M. Dembitzer, N. R. Ghatak, K. J. Fan, H. M. Zimmerman, On the relationship between human and experimental granule cell type cerebellar degeneration. Journal of neuropathology and experimental neurology 32, 493–502 (1973).

85. B. Alvarez-Castelao, E. M. Schuman, The Regulation of Synaptic Protein Turnover. J Biol Chem 290, 28623–28630 (2015).

86. C. Fruhbeis, D. Frohlich, E. M. Kramer-Albers, Emerging roles of exosomes in neuron-glia communication. Front Physiol 3, 119 (2012).

87. S. Saman et al., Exosome-associated tau is secreted in tauopathy models and is selectively phosphorylated in cerebrospinal fluid in early Alzheimer disease. J Biol Chem 287, 3842–3849 (2012).

88. K. Yuyama, H. Sun, S. Mitsutake, Y. Igarashi, Sphingolipid-modulated exosome secretion promotes clearance of amyloid-beta by microglia. J Biol Chem 287, 10977–10989 (2012).

89. Y. Iguchi et al., Exosome secretion is a key pathway for clearance of pathological TDP-43. Brain 139, 3187–3201 (2016).

90. R. I. Morimoto, Proteotoxic stress and inducible chaperone networks in neurodegenerative disease and aging. Genes Dev 22, 1427–1438 (2008).

91. A. Di Fonzo et al, FBXO7 mutations cause autosomal recessive, early-onset parkinsonian-pyramidal syndrome. Neurology 72, 240–245 (2009).

92. C. Paisan-Ruiz et al., Early-onset L-dopa-responsive parkinsonism with pyramidal signs due to ATP13A2, PLA2G6, FBXO7 and spatacsin mutations. Movement disorders: official journal of the Movement Disorder Society 25, 1791–1800 (2010).

93. X. Li, D. Thompson, B. Kumar, G. N. DeMartino, Molecular and cellular roles of PI31 (PSMF1) protein in regulation of proteasome function. The Journal of biological chemistiy 289,17392–17405 (2014).

94. J. M. Deger, J. E. Gerson, R. Kayed, The interrelationship of proteasome impairment and oligomeric intermediates in neurodegeneration. Aging Cell 14, 715–724 (2015).

95. K. Tanaka, N. Matsuda, Proteostasis and neurodegeneration: the roles of proteasomal degradation and autophagy. Biochim Biophys Acta 1843, 197–204 (2014).

96. D. Vilchez, I. Saez, A. Dillin, The role of protein clearance mechanisms in organismal ageing and age-related diseases. Nat Commun 5, 5659 (2014).

97. F. J. Dennissen, N. Kholod, F. W. van Leeuwen, The ubiquitin proteasome system in neurodegenerative diseases: culprit, accomplice or victim? Prog Neurobiol 96,190–207 (2012).

98. S. J. Guyenet et al., A simple composite phenotype scoring system for evaluating mouse models of cerebellar ataxia. J Vis Exp 10.3791/1787 (2010).

99. R. J. Carter et al., Characterization of progressive motor deficits in mice transgenic for the human Huntington’s disease mutation. J Neurosci 19, 3248–3257 (1999).

100. M. S. Brill et al., Branch-Specific Microtubule Destabilization Mediates Axon Branch Loss during Neuromuscular Synapse Elimination. Neuron 92, 845–856(2016).

