## Supplemental Figures for "The proteasome regulator PI31 is required for protein homeostasis, synapse maintenance and neuronal survival in mice"

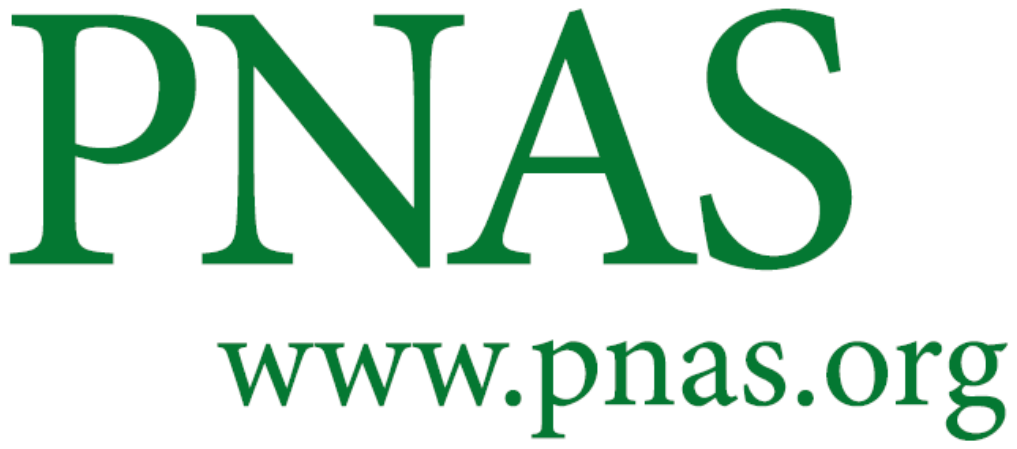

Supplementary Information for

**The proteasome regulator PI31 is required for protein homeostasis, synapse maintenance and neuronal survival in mice**

Adi Minis^1,*^, Jose Rodriguez^1,*^, Avi Levin^1,3*^, Kai Liu^1^, Eve-Ellen Govek^2^, Mary E. Hatten^2^, Hermann Steller^1^

Mary E. Hatten

**This PDF file includes:**

Figures S1 to S4

Legends for Movies S1 to S3

**Other supplementary materials for this manuscript include the following:**

Movies S1 to S3

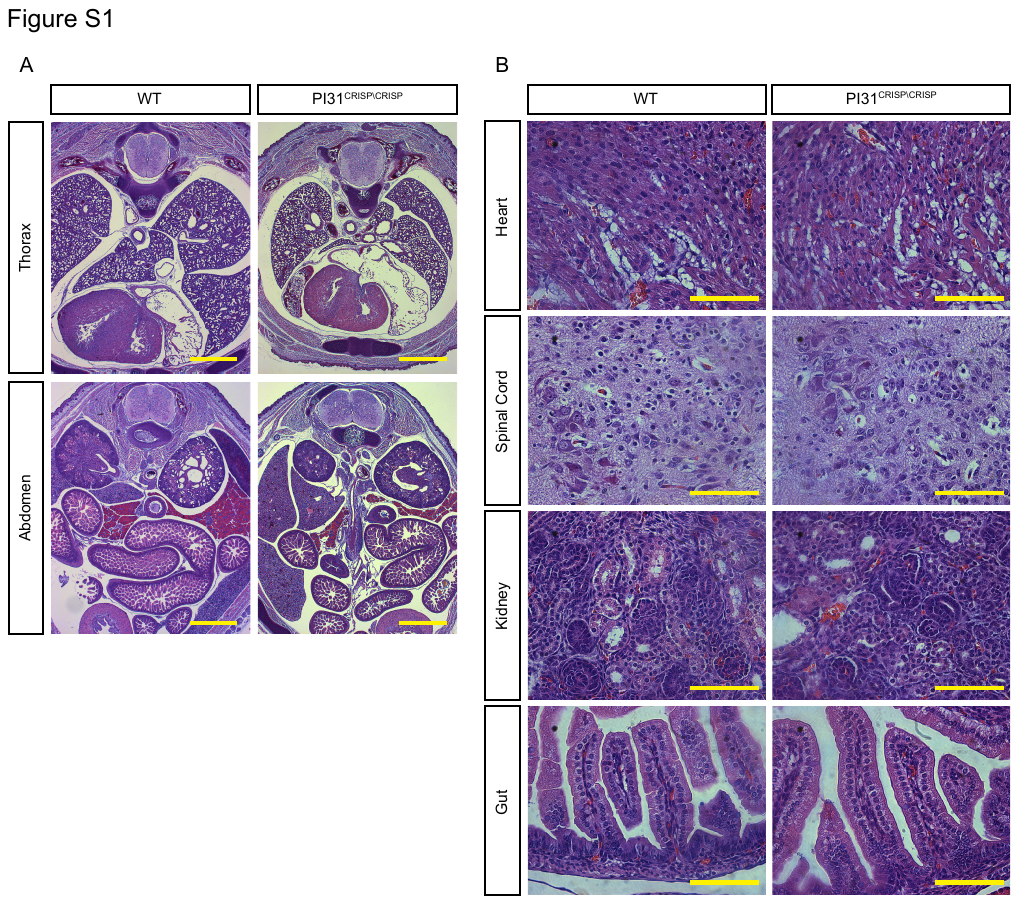

**Fig. S1**. Histological analysis of E18.5 PI31^CRISP\CRISP^ embryos. Low (A) and high (B) magnification of H&E coronal sections taken from different levels of the apical-caudal axis of E18.5 PI31^CRISP\CRISP^ and control littermates. Loss of PI31 did not have profound consequences for the development of many embryonic tissues and organs. Scale bars are 1mm and 0.01mm for figures in panel A and B respectively.

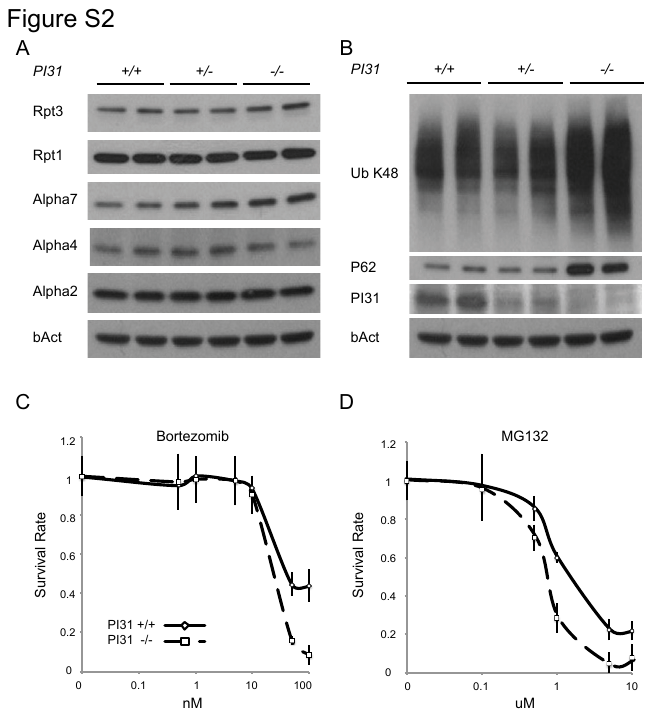

**Fig. S2.** **Loss of PI31 sensitizes mouse embryonic fibroblasts to proteotoxic stress (***A*) Analysis of proteasomal proteins in PI31-Null MEFs. Western blot analysis of WT, heterozygous and PI31 KOF MEFs for individual proteasome subunits does not indicate substantial changes. (*B*) Western blot analysis of WT, Heterozygous and KOF MEFs shows accumulation of poly-ubiquinated proteins (Ub K48) and increased levels of the stress marker p62 in KO MEFs. C, D Diagrams showing survival rate of WT and KOF MEFs treated with Bortezomib (*C*) and MG 132 (*D*) indicated concentrations (average of 3 experiments, each experiment was done in triplicate).

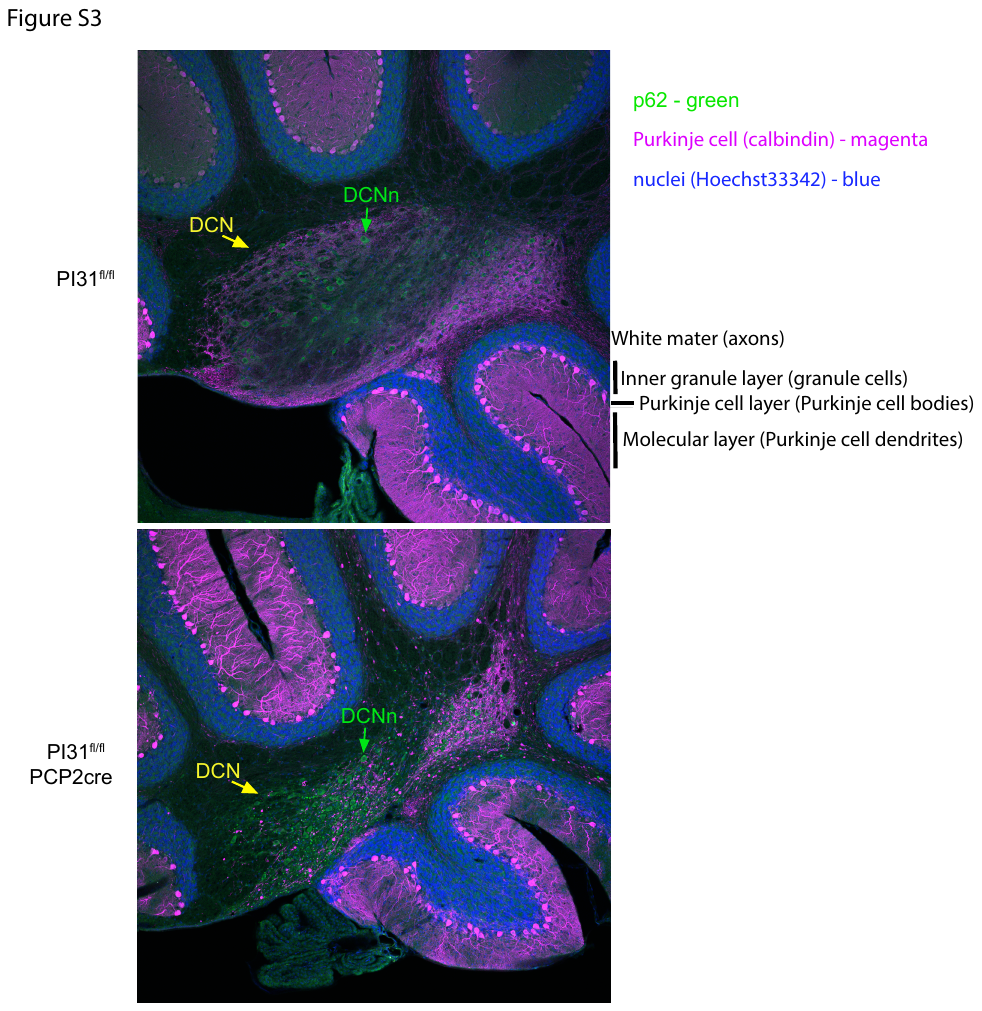

**Fig. S3.** p62-positive aggregates (green) accumulated in the DCN (yellow arrow) of 2-month old *PI31^fl/fl^ PCP2-cre* mice. The size of the DCN was reduced in *PI31^fl/fl^ PCP2-cre* mice, presumably due to loss of PC axon terminals (magenta).

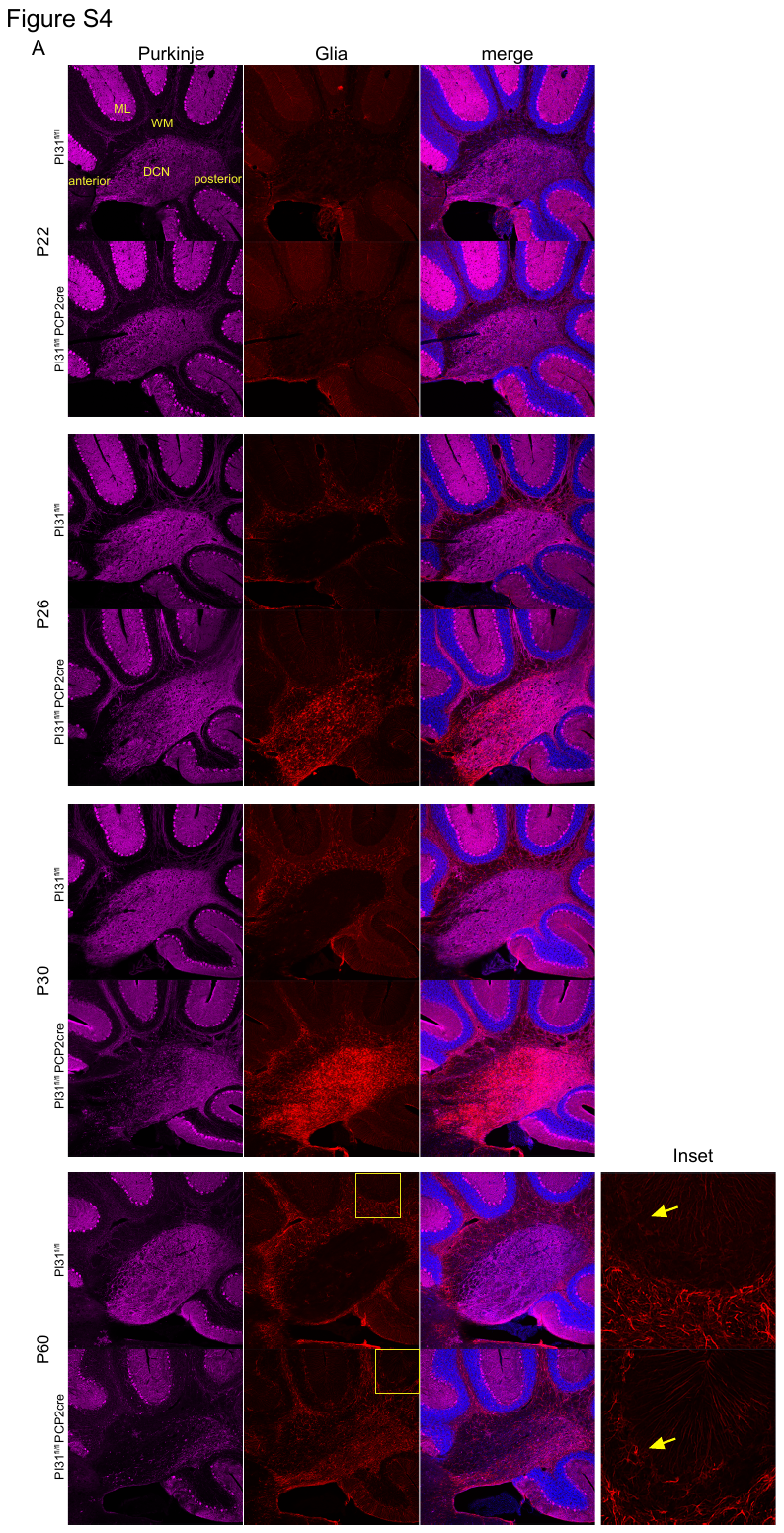

**Fig. S4.** **Loss of PI31 in Purkinje cells leads to gliosis in the DCN of the cerebellum**. Cerebellum from P22, P26, P30 and P60 *PI31^fl/fl^* and *PI31^fl/fl^ PCP2-cre* mice stained with Calbindin (magenta) for Purkinje cells, GFAP (red) for glia, and Hoechst 33342 for nuclei (blue). PC axon terminals in the DCN of *PI31^fl/fl^ PCP2-cre* looked normal at P22, but evidence of axon swelling and loss became apparent at P26 and prominent at P30. At P60, loss of axon terminals in *PI31^fl/fl^ PCP2-cre* mice was nearly complete. GFAP staining in cerebellum of *PI31^fl/fl^ PCP2-cre* mice was similar to that of control littermates at P22. At P26, gliosis in the DCN became apparent, and at P30 there was a dramatic increase of glia staining strongly for GFAP in the DCN. This reaction appears to be specific to the PC axons, as it was not observed in the Purkinje or molecular layers. At P60, there were also some reactive astrocytes in the Purkinje layer (inset), coinciding with the onset of PC loss. DCN (deep cerebellar nuclei), ML (molecular layer) and WM (white matter). Scale bar 100 µm.

**Movie S1 (separate file).** **Illustrating neuro-motor phenotype of PI31^fl/fl^ Cdx2-cre mice**

14 days old *PI31^fl/fl^* and *PI31^fl/fl^ CDX2-cre* littermates. *PI31^fl/fl^ CDX2-cre* mice had sever neuromotor phenotypes, characterized by spasticity, rigid muscle tone, strong tremor and a severely impaired righting response.

**Movie S2 (separate file).** **Illustrating motor defects of PI31^fl/fl^ Hb9-cre mice**

5 month-old *PI31^fl/fl^* and *PI31^fl/fl^ Hb9-cre* littermates. *PI31^fl/fl^ Hb9-cre* mouse had kyphosis and breathing difficulties.

**Movie S3 (separate file). Illustrating cerebellar behavioral defects in *PI31^fl/fl^ PCP2-cre* mice**

Movie of 7 month-old male *PI31^fl/fl^* and *PI31^fl/fl^ PCP2-cre* siblings. The *PI31^fl/fl^ PCP2-cre* mouse had a halting gait and lost balance frequently, falling back or to the left or right. *PI31^fl/fl^ PCP2-cre* mouse has difficulty regaining balance as it falls to the left.
